# Versatile simulations of admixture and accurate local ancestry inference with *mixnmatch* and *ancestryinfer*

**DOI:** 10.1101/860924

**Authors:** Molly Schumer, Daniel L. Powell, Russ Corbett-Detig

**Affiliations:** Department of Biology, Stanford University; Centro de Investigaciones Científicas de las Huastecas “Aguazarca”; Hanna H. Gray Fellow, Howard Hughes Medical Institute; Department of Biology, Texas A&M University; Genomics Institute, University of California, Santa Cruz; Department of Biomolecular Engineering, University of California, Santa Cruz

**Keywords:** hybridization, admixture, local ancestry inference, hidden Markov model

## Abstract

It is now clear that hybridization between species is much more common than previously recognized. As a result, we now know that the genomes of many modern species, including our own, are a patchwork of regions derived from past hybridization events. Increasingly researchers are interested in disentangling which regions of the genome originated from each parental species using local ancestry inference methods. Due to the diverse effects of admixture, this interest is shared across disparate fields, from human genetics to research in ecology and evolutionary biology. However, local ancestry inference methods are sensitive to a range of biological and technical parameters which can impact accuracy. Here we present paired simulation and ancestry inference pipelines, *mixnmatch* and *ancestryinfer*, to help researchers plan and execute local ancestry inference studies. *mixnmatch* can simulate arbitrarily complex demographic histories in the parental and hybrid populations, selection on hybrids, and technical variables such as coverage and contamination. *ancestryinfer* takes as input sequencing reads from simulated or real individuals, and implements an efficient local ancestry inference pipeline. We perform a series of simulations with *mixnmatch* to pinpoint factors that influence accuracy in local ancestry inference and highlight useful features of the two pipelines. Together, *mixnmatch* and *ancestryinfer* are powerful tools for predicting the performance of local ancestry inference methods on real data.

## Introduction

Since the advent of inexpensive whole genome sequencing, it has become increasingly clear that hybridization is an important part of the evolutionary history of many species. This has made methods to study hybridization fundamental tools in the fields of genetics and evolutionary biology. In addition to methods for inferring the genome-wide history of admixture (Alexander, Novembre, & Lange, 2009; Patterson et al., 2012; Pritchard, Stephens, & Donnelly, 2000), researchers have recently taken advantage of methods that make it possible to infer ancestry at a small spatial scale along the genome (i.e. “local ancestry inference”). Originally applied to study genetic diseases in humans (Hoggart, Shriver, Kittles, Clayton, & McKeigue, 2004; Montana & Pritchard, 2004; Patterson et al., 2004; Winkler, Nelson, & Smith, 2010), applications of local ancestry inference methods have become a cornerstone of studies of genome evolution (Sankararaman et al., 2014; Schumer et al., 2018), population history (Baharian et al., 2016; Corbett-Detig & Nielsen, 2017), and trait evolution (Heliconius Genome, 2012; Jones et al., 2018; Oziolor et al., 2019).

Particularly popular among local ancestry inference methods are methods that use a hidden Markov model (HMM) to infer ancestry as a hidden state based on observations of genotypes or sequencing read counts. For autosomal loci in diploid individuals, there are three possible ancestry states (homozygous parent species 1, homozygous parent species 2, and heterozygous for the two ancestries). Ancestry HMMs allow the probability of a given local ancestry state to be modeled as a function of the observed data in the region, the ancestry state probability at the previous site, and the recombination distance between adjacent ancestry informative sites, among other possible parameters. The output of these methods is typically posterior probabilities for each possible ancestry state at each ancestry informative site along the chromosome (Alexander et al., 2009; Andolfatto et al., 2011; Corbett-Detig & Nielsen, 2017).

As local ancestry methods have been applied in more and more species (e.g. Cande, Andolfatto, Prud’homme, Stern, & Gompel, 2012; R. Li et al., 2018; Sankararaman et al., 2014; Schumer et al., 2014; Slotte et al., 2013), each with their own population genetic properties and demographic histories, simulation tools to evaluate performance have not kept apace. While some studies have carefully modeled the demographic history of hybridizing populations and the impact of this history on the accuracy of ancestry inference (e.g. Sankararaman et al., 2014; Medina, Thornlow, Nielsen, & Corbett-Detig, 2018), this typically requires the development of custom computational pipelines. As a result, many studies do not evaluate the expected performance of local ancestry inference methods and use default parameter sets.

While the majority of tools for local ancestry inference report performance under parameters relevant to human populations (e.g. Maples, Gravel, Kenny, & Bustamante, 2013), these tools are being applied much more broadly. This is concerning because local ancestry inference approaches that perform well in one context may perform poorly in others, as their performance is sensitive to a number of biological and technical variables (Medina et al., 2018; Sankararaman et al., 2014; Schumer, Cui, Rosenthal, & Andolfatto, 2016). Because of the importance of accurate local ancestry inference in evolutionary biology and genetics, simulation tools that allow researchers to systematically evaluate their accuracy and common biases are needed.

Here, we present a hybrid genome simulation pipeline called *mixnmatch* that can be used to evaluate the accuracy of local ancestry inference under a range of biological and technical parameters. Our pipeline builds on previous simulation tools developed by us and others to implement flexible demographic simulations in both ancestral parental populations and hybrid populations. Species-specific genetic parameters, including base composition and local recombination rates, can be incorporated into simulations. Users also specify a number of technical parameters that impact accuracy, including sequencing depth, sequencing error, and cross-contamination rates.

In addition to simulating admixed genomes with *mixnmatch*, we provide a paired pipeline called *ancestryinfer* that implements local ancestry inference on real or simulated data. This pipeline builds off of a previously published HMM (AncestryHMM; Corbett-Detig & Nielsen, 2017) and includes several features that seamlessly integrate its use with raw sequence data, automating and parallelizing steps from mapping Illumina reads to the output of posterior probabilities for ancestry states. Moreover, both pipelines are user-friendly, with parameters defined in a text-editable configuration file and automated and docker-based installation options. Together, *mixnmatch* and *ancestryinfer* will make it feasible for users to perform sophisticated simulations to predict accuracy and apply the same approaches when analyzing their data.

## Methods & Results

### Overview

The overall structure of *mixnmatch* is described in Figure 1 and in more detail in Figure S1. First, the pipeline simulates parental haplotypes, either using user-provided genomes or the coalescent simulator *macs* (Chen, Marjoram, & Wall, 2009), which allows for simulations of complex population history (Figure S1; Supporting Information 1). With *macs*-based simulations, users can still provide one of the parental genomes for use as a base sequence (Figure S1).

**Figure 1.**
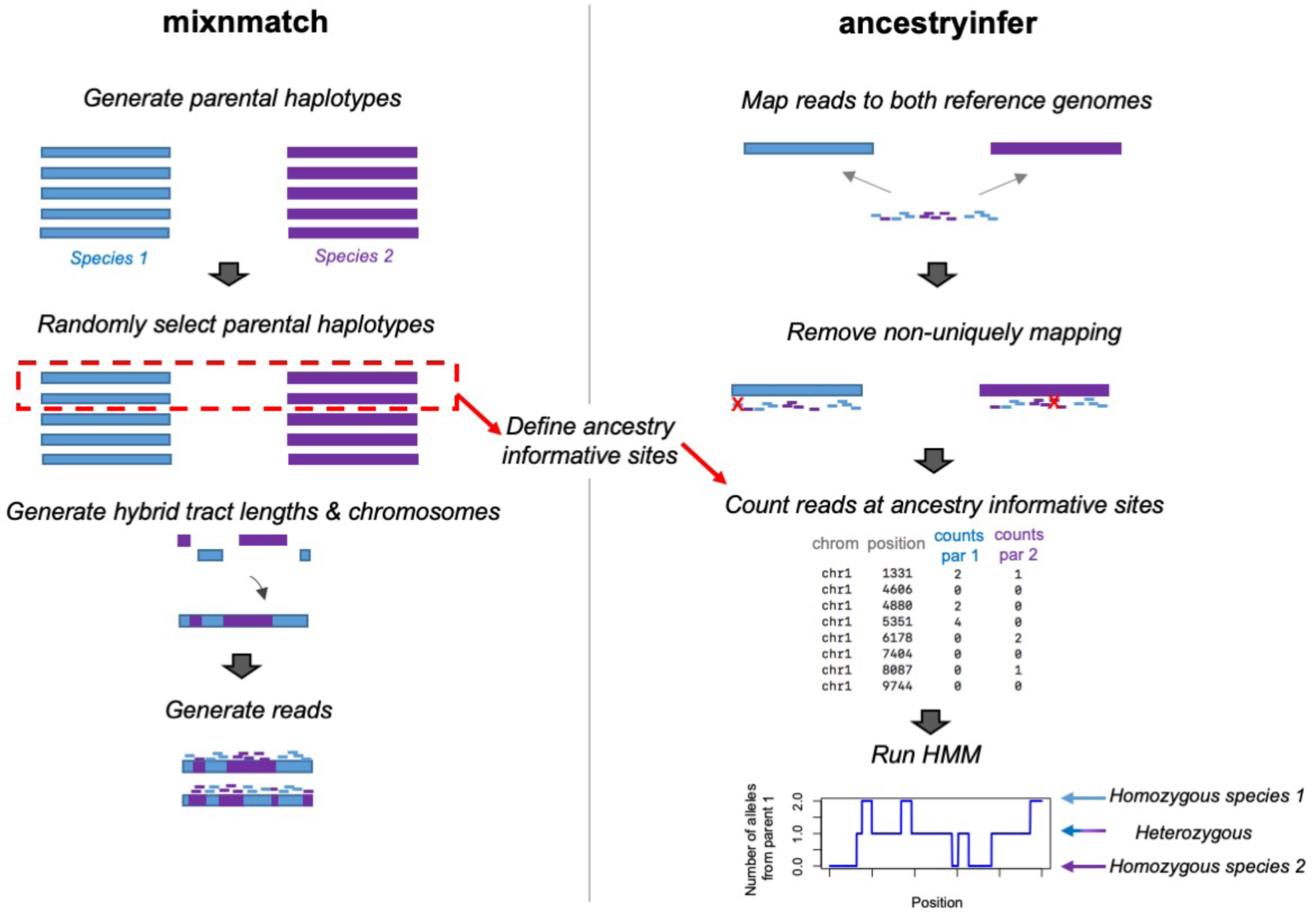
Schematic showing the major steps in the *mixnmatch* (left) and *ancestryinfer* (right) pipelines. *mixnmatch* can be used to simulate hybrid data under user-specified parental and hybrid demographic scenarios and under a range of technical parameters. A more detailed description of the *mixnmatch* pipeline and its options is outlined in Figure S1. Simulated Illumina data output by *mixnmatch* can be input into our automated ancestry inference pipeline, *ancestryinfer*, as can data from natural and artificial hybrids. The red box indicates the haplotypes chosen from the parental species during *mixnmatch* to identify ancestry informative sites that can later be used in *ancestryinfer*. Blue shading indicates tracts and reads derived from parent species 1 and purple indicates tracts and reads derived from parent species 2. Red X’s mark reads that will be removed for not mapping uniquely to both parental genomes.

Next, ancestry tract lengths for hybrid individuals are generated with SELAM (Corbett-Detig & Jones, 2016), using information about the admixture proportion, demographic history of the hybrid population, and number of generations since initial admixture. Based on these tract lengths, hybrid genomes are constructed from previously simulated parental haplotypes. For each individual, reads are generated at user-specified lengths and depth using the *wgsim* program (H. Li, 2011). If desired, users can simulate cross-contamination between samples during read generation. Together, these features of *mixnmatch* allow for simulation of experimental design choices and the history of both the parental and hybrid population populations, making it a powerful tool for studies of hybridization.

Output files produced by the *mixnmatch* simulator include fastq and fasta files for each individual, a bed file indicating true ancestry along the simulated chromosome, and all necessary files for running an efficient local ancestry inference HMM, implemented through our associated pipeline, *ancestryinfer* (Table S1). After running local ancestry inference with *ancestryinfer*, users can evaluate the performance of local ancestry inference with a provided script that summarizes accuracy by comparing inferred versus true ancestry at each ancestry informative site.

### Installation

Users can install the two pipelines using step-by-step instructions provided with *mixnmatch* and *ancestryinfer* or by loading a docker image with all dependencies for both pipelines pre-installed (see Appendix 1; user manual). Parameters for each pipeline are set in a text-editable configuration file (examples available on github: https://github.com/Schumerlab/mixnmatch; https://github.com/Schumerlab/ancestryinfer). Instructions for setting parameters can be found in the *mixnmatch* and *ancestryinfer* user manuals (see Appendix 1-2). *mixnmatch* and *ancestryinfer* can be parallelized with a SLURM resource management system (Appendix 1-2) and the non-parallel version can be run any Linux system or on Mac operating systems.

### Simulation of parental haplotypes and definition of ancestry informative sites

Generating parental haplotypes is the first step in *mixnmatch* simulations (Figure S1). An important feature of *mixnmatch* is the ability to simulate demographic history. This is crucial because the population history of each parental species will influence genetic diversity and the extent of background linkage disequilibrium within species, as well as divergence between species, all of which can impact accuracy in local ancestry inference. To model this, users provide a *macs* command (Chen, Marjoram, & Wall, 2009) describing the demographic history of the two parental species in the configuration file. *mixnmatch* executes this command and converts the *macs* output to nucleotide sequences using the *seq-gen* program (Rambaut & Grassly, 1997). Users can optionally provide species-specific base composition and transition/transversion ratios in the configuration file, as well as a local recombination map. If provided, this recombination map will be used in the *macs* simulations of parental haplotypes and in generating hybrid ancestry tracts (see *Simulation of hybrid genomes*).

One of these simulated sequences from each parental population is set aside as to be used as the reference sequence. The remaining haplotypes are then used to define ancestry informative markers and are later used to generate hybrid chromosomes (see *Simulation of hybrid genomes* below). Ancestry informative markers are defined as highly differentiated markers among a randomly selected subset of the simulated parental haplotypes. Users specify the required frequency difference between species and the number of parental haplotypes to use to evaluate this in the configuration file. We note that the appropriate choices for these values will depend on the level of divergence and shared polymorphisms between species; users can rely on predictions from population genetic theory (e.g. Wakeley & Hey, 1997) or *mixnmatch* simulations to explore these parameters. Together, these steps model the impacts of demographic history on the number and distribution of ancestry informative sites, as well as the process researchers typically follow in identifying them.

#### Other options for parental haplotype generation

Although the method of haplotype generation described above incorporates demography, recombination, and incomplete lineage sorting in the parental species, it lacks other complexities of real genome sequences such as repetitive elements and local variation in base composition. To accommodate these additional challenges, we allow users to provide an ancestral sequence to which simulated mutations are added. This option incorporates features of real sequences while modeling mutations and ancestral recombination events using a coalescent framework.

Another biological variable that can impact accuracy in ancestry HMMs is drift between the reference panel used to define ancestry informative sites and the populations that actually contributed to the hybridization event. Users can tell *mixnmatch* to allow drift between the source population and the population used to define ancestry informative sites (using *macs*). If users choose this option, ancestry informative sites are defined based on the drifted parental population *instead* of the hybridizing parental population. This generates realistic allele frequency differences between the reference and source parental populations, as well as covariance in allele frequency differences due to linkage. In real data, this could contribute to errors in downstream analysis.

We also provide an option for using *mixnmatch* with the exact reference genomes and ancestry informative sites that users plan to rely upon in their experiments. Instead of a *macs* command describing parental population history, this option takes the two parental reference genomes and a number of population genetic parameters as input (Figure S1). This approach is described in Supporting Information 1 (see also Schumer et al., 2016). Although this option has some limitations that should be considered (Supporting Information 1), it allows users to evaluate performance on species-specific reference genomes and ancestry informative sites.

### Simulation of hybrid genomes

In addition to allowing users to simulate the demographic history of the parental species, *mixnmatch* models the demographic history of the hybrid population (Figure S1). Any process that influences the distribution of ancestry tract lengths, from bottlenecks to assortative mating, could impact the accuracy of local ancestry inference. To incorporate this, *mixmatch* uses a previously developed tool (SELAM, Corbett-Detig & Jones, 2016) to model the effects of demography on ancestry tracts. Users specify the mixture proportions contributed from each parent species to the hybrid population and the number of generations since initial admixture in the *mixnmatch* configuration file. In addition, users can choose to provide a parameter file describing the demographic history of the hybrid population (Appendix 1) as well as selection on hybrids.

Using these tract lengths, *mixnmatch* next generates hybrid chromosomes. For each ancestry tract, *mixnmatch* extracts the focal region from a randomly selected parental haplotype of the appropriate ancestry. If users have provided a local recombination map, *mixnmatch* uses this in converting tract coordinates from genetic to physical distance; otherwise a global recombination rate is used. This process is repeated until an entire haplotype is generated, and two such haplotypes are combined to generate both chromosomes within a diploid hybrid individual. Importantly, this approach introduces variation to the simulation from processes such as incomplete lineage and sampling of a reference panel, both of which can impact downstream accuracy.

The pipeline next simulates reads uniformly from these hybrid chromosomes using the *wgsim* program (H. Li, 2011), with user specified read lengths, read mate type, coverage, indel and error rates. At the same time contamination can be simulated. During this step *mixnmatch* writes out the true ancestry for each individual at every position along the chromosome, facilitating analysis of accuracy downstream.

The final output of *mixnmatch* includes all of the files needed for running our paired pipeline for local ancestry inference, *ancestryinfer*, as well as files needed for other local ancestry inference tools. These include simulated reference genomes, ancestry informative sites and counts for each allele in the parental reference panel, simulated Illumina reads, and bed formatted files containing the true ancestry for each individual (Table S1).

One possible shortcoming of our approach for generating hybrid haplotypes is that it does not model coalescence among samples after admixture, which could generate errors in local ancestry inference not captured by our simulation approach. This is most likely to impact simulations of very small populations or ancient admixture (Corbett-Detig & Nielsen, 2017). We also note that the number of parental haplotypes used to generate the hybrid chromosomes is determined by the total number of parental haplotypes users choose to simulate. Simulating fewer parental haplotypes will decrease *mixnmatch* runtime, but users should ensure that the total number of parental haplotypes simulated captures most of the genetic variation within the parental populations (e.g. Figure S2; Watterson, 1975).

### A versatile ancestry inference pipeline

To facilitate local ancestry inference analysis of real and simulated data, we developed a paired pipeline called *ancestryinfer*. This pipeline automates steps from read mapping to local ancestry inference, and is easy-to-use and parallelizable (Supporting Information 2). Briefly, the work flow of this pipeline (Figure 1) begins with mapping reads from a hybrid individual to both parental references independently with *bwa mem* (H. Li & Durbin, 2009) and identifying reads that do not map uniquely to either of the parental genomes. These reads are then excluded from the hybrid individual’s bam file using *ngsutils* (Breese & Liu, 2013). Such reads may fall within repetitive regions of the parental genomes, be impacted by mapping bias, incompleteness of one parental reference, or insertions/deletions that disrupt mapping. These technical issues have received less attention as it relates to their impact on local ancestry inference (Supporting Information 3) but have been shown to have major impacts in other types of analyses such as allele-specific expression (Degner et al., 2009; Stevenson, Coolon, & Wittkopp, 2013).

Next, reads matching each parental allele at ancestry informative sites are counted from a *samtools* mpileup file (H. Li, 2011) generated for each hybrid individual. There are two options in the pipeline for identifying ancestry informative sites. If the genomes provided by the user are co-linear, users can direct *ancestryinfer* to automatically identify sites that differ between them. Alternately, users can provide the locations of ancestry informative sites and their estimated frequencies in the parental species. The latter option allows users to take advantage of reference assemblies for both species if they are available.

Counts for each parental allele at ancestry informative sites are subsampled to thin to one ancestry informative site per read if multiple sites occur within one read. We implement this thinning because mismapping can generate clusters of errors and non-independence between sites is not modeled in the HMM. Finally, Ancestry_HMM (Corbett-Detig & Nielsen, 2017) is applied to infer posterior probabilities of each ancestry state at ancestry informative sites along the genome. In addition, *ancestryinfer* summarizes the intervals over which ancestry transitions occur (Figure S3). With the exception of false switches in ancestry generated by errors, these intervals reflect observed crossover events in hybrids. These crossover intervals can be used to generate recombination maps (see below, *Generating a hybrid recombination map using observed ancestry transitions*).

If users have run the *ancestryinfer* pipeline on data simulated by *mixnmatch*, the accuracy of local ancestry inference can be summarized by running a script provided with the *mixnmatch* pipeline (Appendix 2). Briefly, the script generates hard-calls at a user-specific posterior probability threshold and compares true and inferred ancestry at each ancestry informative site along the chromosome. The output of this script includes plots summarizing individual-level accuracy, accuracy as a function of tract length (Figure S4), and a file tabulating all accurate and inaccurate calls in individual tracts as well as mean posterior error.

### Predicted accuracy of local ancestry inference in simulated data

#### Basic simulation setup

Using *mixnmatch* and *ancestryinfer*, we next tested the accuracy of local ancestry inference with simulated data under a range of biological and technical scenarios. For these simulations, we started with a base parameter set (Table S2) and then systematically modified parameters in individual simulations. For this base parameter set we simulated 200 generations since initial admixture, 50-50 mixture proportions between the two parental species, per site polymorphism rates in each of the parental species of 0.1%, and pairwise sequence divergence between the parental species of 0.5%. We used the first 10 Mb of chromosome 1 from the swordtail fish species *Xiphophorus birchmanni* as the ancestral sequence, and provided *mixnmatch* with an inferred recombination map (Schumer et al., 2018) from that same region.

We simulated 100 parental haplotypes for each species, which will capture the majority of parental polymorphisms segregating in these populations (Figure S2; Watterson, 1975). We sampled 20 parental haplotypes from both species to define ancestry informative sites and required a frequency difference of 95% between species for a site to be treated as ancestry informative. A complete description of these simulations can be found in Supporting Information 4. For this set of parameters, the pipeline required a total of 0.66 CPU hours and was run with 96 Gb of memory for the sequence generation step and 64 Gb of memory for the hybrid genome simulation step. Simulations were performed on Dell C6420 servers on Stanford’s Sherlock High Performance Computing cluster.

In simulations with this parameter set, accuracy of local ancestry inference was high, with estimated error rates of <0.5% per ancestry informative site (Figure S4). As expected, shorter ancestry tracts have higher per-basepair error rates (Figure S4). Because tract lengths often differ systematically across the genome and in particular around selected sites (Sedghifar, Brandvain, Ralph, & Coop, 2015; Shchur, Svedberg, Medina, Corbett-Detig, & Nielsen, 2019), this highlights the importance of considering local error rates throughout the genome when analyzing local ancestry data (Supporting Information 4, Figure S5).

#### Simulations under a range of scenarios

To understand what biological and technical variables impact accuracy in local ancestry inference, we next modified individual parameters in turn. Below we summarize the scenarios we tested that had the greatest impact on accuracy. A full description of our simulations can be found in Supporting Information 4.

Intuitively, with increasing divergence between species, there will be more ancestry informative sites. This is predicted to result in more accurate local ancestry inference. To evaluate this, we performed simulations varying pairwise divergence between the hybridizing species from 0.25% to 1%. We note that these simulations focus on deeper divergence than what has been considered in previous work (Maples et al., 2013; Medina et al., 2018) but span realistic levels of divergence found in naturally hybridizing species (Brandvain, Kenney, Flagel, Coop, & Sweigart, 2014; Schumer et al., 2014; Teeter et al., 2008; Turissini & Matute, 2017). As expected, accuracy increased with higher divergence between the hybridizing species (Figure 2), as did the precision with which the locations of ancestry transitions were identified (Figure S3; consistent with previous results, Medina et al., 2018).

**Figure 2.**
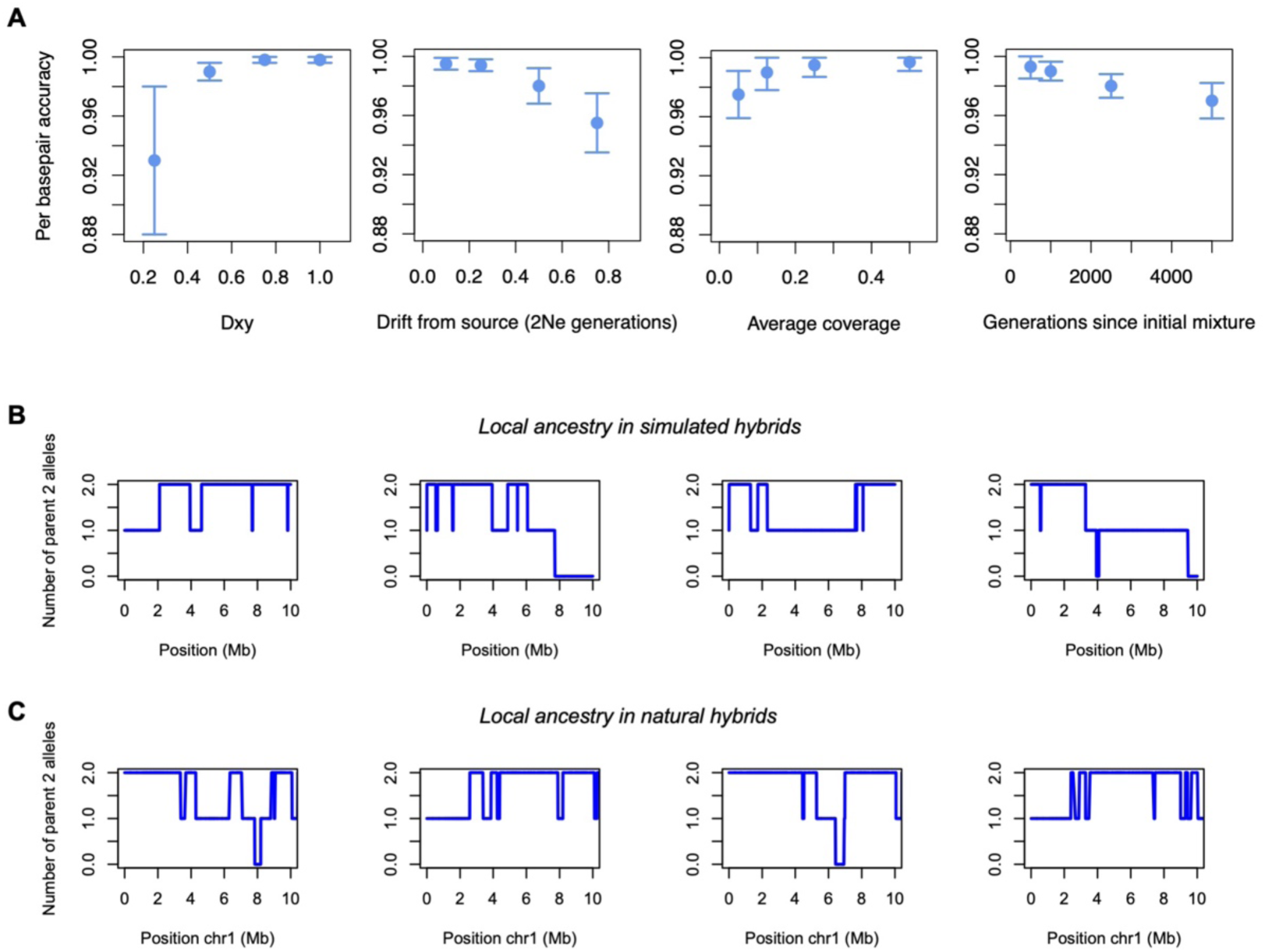
A) Results of *mixnmatch* simulations evaluating accuracy under different biological and technical parameters. All simulations start with the same basic parameter set (Table S2) and systematically vary the focal parameter (see Supporting Information 2). Points indicate the mean of individual-level accuracy and whiskers indicate two standard deviations. B) Example local ancestry results for simulated hybrids. Parameter used in this simulation were ∼1X average coverage, sequence divergence of 0.5%, within species polymorphism rates of 0.1%, 20 generations since initial admixture, and 35% of the genome derived from parent species 1. C) Example local ancestry results for the first 10 Mb of chromosome 1 for *X. birchmanni* x *X. malinche* hybrid individuals from the Chahuaco falls hybrid population, where per-base coverage and inferred parameter values match those simulated in B. Note the qualitative similarities between C and B in the number of ancestry transitions and the size of the ancestry tracts. In C parent 2 alleles are those derived from the *X. malinche* parental species.

As the time since initial admixture increases, recombination events in each generation split haplotypes of a given ancestry into smaller and smaller pieces. Since these short tracts will contain fewer ancestry informative sites, this leads to the prediction that local ancestry inference will be more accurate in populations that have hybridized recently, which is indeed what we observed (Figure 2).

Following a similar logic, skewed admixture proportions are expected to reduce accuracy of ancestry inference in tracts derived from the “minor” parental species (i.e. the parental species that contributed less to the initial hybridization event). This is because only recombination events that occur in regions heterozygous for ancestry from the two parent species are detectable with ancestry HMMs, and minor parent haplotypes are more frequently found in this state (Gravel, 2012). As expected, we observe that the accuracy of local ancestry inference within minor parent tracts is reduced in simulations with skewed initial admixture proportions (Figure S6).

Ideally, reference panels for defining ancestry informative sites should be derived from the same parental populations that contributed to the admixture event. In practice, this is often not possible since source populations may no longer exist, may be unknown, or may themselves be admixed, making it more sensible to use allopatric populations for a reference panel. However, such populations are also expected to have some level of genetic drift from the admixing populations, which could impact accuracy. In *mixnmatch* this can be modeled by adding drift to the simulation and specifying which populations to use in defining ancestry informative sites.

To investigate the impact of using reference panels with drift from the hybridizing populations, we used *mixnmatch* to simulate two additional populations that split from the parental source populations before hybridization and treated these populations as the reference panel (0.4-3*Ne* generations ago in different simulations, with initial divergence between species occurring 8*Ne* generations ago; Supporting Information 4). We found that accuracy substantially decreased with increasing drift between the reference population and the source parental populations (Figure 2). Notably, this can be partially remediated by increasing the required frequency difference between the parental populations when defining ancestry informative sites (Supporting Information 4).

A common decision that researchers make is how much coverage to collect per sample. Intuitively, early generation hybrids will require less data to accurately infer local ancestry than later generation hybrids because of differences in the distribution of ancestry tract lengths. We simulated genome-wide coverage between 0.05-0.5X with *mixnmatch* (Supporting information 4). As expected, increased coverage improved accuracy but our results also suggested that beyond a certain level of coverage, improvements in accuracy plateau (Figure 2). However, higher coverage continued to improve the resolution of the locations of ancestry transitions (Figure S7).

In general, we find that the HMM implemented in *ancestryinfer* (Corbett-Detig & Nielsen, 2017) is not particularly sensitive to user-provided priors for admixture time or admixture proportion, but is somewhat sensitive to recombination rate priors (Supporting Information 4). Providing a local recombination prior in *ancestryinfer* modestly increases accuracy when the recombination map does not contain errors (Figure S8). However, in practice recombination maps will contain errors that depend on the method used for map construction among other factors (Supporting Information 4). With moderate levels of error in map inference our simulations suggest that users may benefit from providing a uniform recombination prior (Figure S8).

In recent years there has been substantial interest in the ecological and evolutionary genomics community in restriction site associated sequencing (or RAD-seq; Andrews, Good, Miller, Luikart, & Hohenlohe, 2016; Peterson, Weber, Kay, Fisher, & Hoekstra, 2012; Van Tassell et al., 2008) as a low-cost option for generating genomic data. However, RAD-seq data may be suboptiomal for local ancestry inference applications. This is because overdispersion in the spacing between sampled sites, coverage variation, and genealogical biases generated by variants in restriction enzyme cut-sites introduced by this method could all reduce the accuracy of local ancestry inference. To explore this, we generated reads *in silico* associated with a commonly used enzyme in RAD (*EcoRI*) but otherwise performed simulations as described above (Supporting Information 5). We found that in the case of *ancestryinfer* performance with RAD data is poor (Figure S9), likely due to the reliance on fewer ancestry informative sites for inference (Supporting Information 5).

Finally, although we focus on modeling accuracy under neutral demographic scenarios, *mixnmatch* can also be used to simulate selection on hybrids. The SELAM program that is used to simulate ancestry tract lengths in *mixnmatch* accommodates selection on hybrid populations (Corbett-Detig & Jones, 2016; Figure S10, Supporting Information 6). This allows users to implement versatile selection scenarios in *mixnmatch* (Appendix 1), and use it to explore signatures of selection on hybrids. For example, recent work has described the impact of selection against hybrid incompatibilities on the number and distribution of ancestry junctions (Hvala, Frayer, & Payseur, 2018), the impact of selection on ancestry tract lengths (Sedghifar et al., 2015; Shchur et al., 2019), and the local frequency of haplotypes derived from the minor parental species (Sankararaman et al., 2014; Schumer et al., 2018; Vernot & Akey, 2014). We demonstrate the use of this feature of *mixnmatch* in Supporting Information 6 and Figure S10.

#### Application to natural and artificial swordtail hybrids

To demonstrate the utility of *ancestryinfer* with real data, we applied it to data from F_1_ and F_2_ hybrids generated from crosses between the swordtail fish species *X. birchmanni* and *X. malinche*. We constructed libraries using a tagmentation based library preparation protocol and collected low coverage whole-genome sequence data for these libraries (∼0.2X coverage per individual, Supporting Information 7, Appendix 3). We used previously collected low-coverage sequence data from 60 individuals of each parental species to estimate allele frequencies at ancestry informative sites (Schumer et al., 2018).

Illumina sequencing data from lab-generated hybrids was input into the *ancestryinfer* pipeline to infer local ancestry along the 24 swordtail chromosomes (here 150 x 2 paired-end data was used). We used the appropriate parameters for mixture proportion, generations since mixture, and global recombination rate based on known information about the cross (see details in Supporting Information 7). We converted posterior probabilities for a given ancestry state into hard calls using a threshold of 0.9 and examined local patterns of ancestry in F_1_ and F_2_ hybrids. We also summarized expected ancestry proportions, heterozygosity in ancestry, and the numbers of observed ancestry transitions genome-wide (Figure 3).

**Figure 3.**
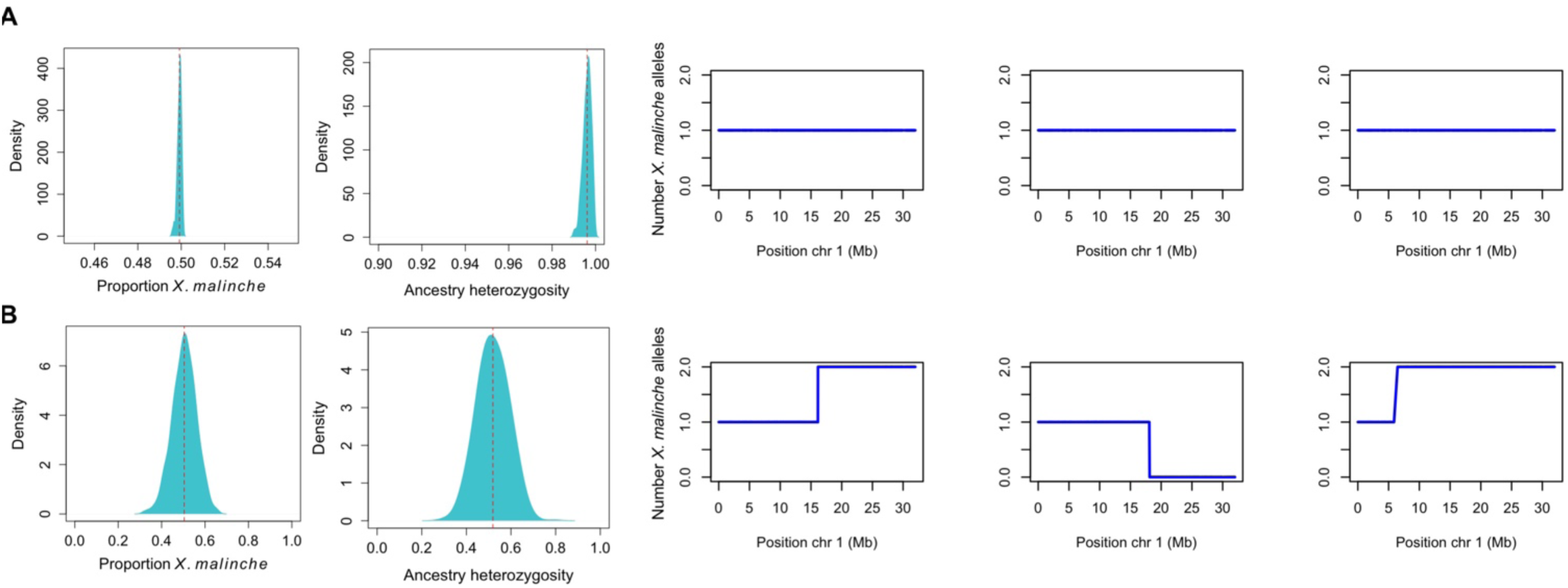
Results of the *ancestryinfer* pipeline run on F_1_ (A) and F_2_ (B) hybrids generated between *X. birchmanni* and *X. malinche*. A) As expected, we infer that F_1_ hybrids have precisely 50% of their genomes derived from each parental species and infer nearly 100% heterozygosity at ancestry informative sites in these individuals. Example local ancestry plots for chromosome 1 for a subset of these F_1_ hybrids are also shown (right). B) Similarly, genome-wide ancestry distributions and genome-wide ancestry heterozygosity in F_2_ hybrids follows predicted distributions. Example local ancestry plots for chromosome 1 for a subset of these F_2_ hybrids are also shown (right).

We find that local and global ancestry patterns in F_1_ and F_2_ hybrids mirror expectations for each cross type (Figure 3), and that the results are consistent with extremely low error rates in ancestry inference. For example, estimated homozygosity at ancestry informative sites in F_1_ hybrids is <0.1% (Figure 3). Importantly, this high level of accuracy is predicted from simulations of early generation hybrids with *mixnmatch* (Supporting Information 7), suggesting that *mixnmatch* simulations are capturing important properties of real data.

#### Generating a hybrid recombination map using observed ancestry transitions

Accurate local ancestry inference has a large number of downstream applications. One such application is inferring the locations of crossovers for the construction of genetic maps (Amores et al., 2014; Rastas, Calboli, Guo, Shikano, & Merilä, 2015; Salomé et al., 2012). As discussed previously, if users specify a posterior probability threshold in the *ancestryinfer* configuration file, the program will output a bed file containing recombination intervals inferred from observed ancestry transitions in hybrids.

We used the locations of observed ancestry transitions in 139 F_2_ hybrids that we generated between *X. birchmanni* and *X. malinche* (at a posterior probability threshold of 0.9; Supporting Information 7-8) to estimate the recombination rate in 5 Mb windows. We used a large window size due to the spatial scale over which ancestry transitions were localized (lower and upper 5% quantile of intervals genome-wide: 23 kb - 667 kb) and because we expected the resulting map to be relatively coarse, given a total of 4038 inferred crossovers genome-wide (average 1.2 per individual per chromosome).

We compared inferred recombination rates in this F_2_ map to a linkage disequilibrium based recombination map for *X. birchmanni* that we had previously generated (Schumer et al., 2018). As expected, we observed a strong correlation in estimated recombination rate between the linkage disequilibrium based and crossover maps (R=0.82, Figure 4, Supporting Information 8). Simulations suggest that the observed correlation is consistent with the two recombination maps being indistinguishable, given the low resolution of the F_2_ map (Supporting Information 8).

**Figure 4.**
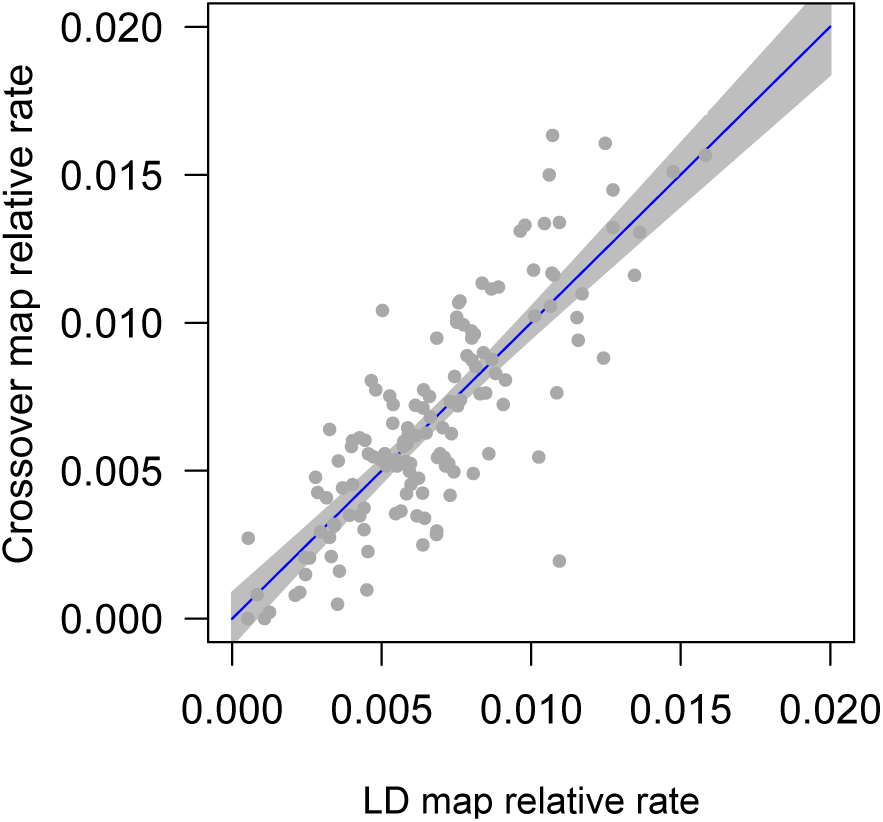
Comparison of F_2_ crossover recombination map generated with *ancestryinfer* and previously published linkage disequilibrium map from the *X. birchmanni* parental species. Relative rates per 5 Mb window for both the linkage disequilibrium map (x-axis) and F_2_ map (y-axis) are shown by gray dots. The blue line shows the best fit regression line between the maps (R^2^ = 0.67) and the gray area shows the 95% confidence intervals. Simulations suggest that the observed correlation is consistent with recombination rates being identical across the two maps (Supporting Information 8).

## Discussion

With an increasing appreciation that hybridization is a common evolutionary process, there has been renewed interest in local ancestry inference in the fields of genetics and evolutionary biology. Accurate local ancestry information is important for applications from admixture mapping to studying genome evolution after hybridization. Despite this, there are few simulation tools that have been developed to model the impacts of biological and technical variables on the accuracy of local ancestry inference.

We demonstrated the use of *mixnmatch* as a flexible tool to predict the accuracy of local ancestry inference under a range of biological scenarios. As expected *a prori*, our simulations show that the factors with the strongest impact on accuracy include the number of ancestry informative sites that distinguish the hybridizing species, the length of the ancestry tracts containing these sites, and the frequency at which sites are erroneously defined as ancestry informative (either due to genetic drift or high levels of shared polymorphisms; Figure 2). We show how *mixnmatch* can also help users make important decisions about their projects, such as how much coverage to collect per hybrid individual and how many parental individuals to sequence to define ancestry informative sites.

*mixnmatch* is primarily designed to allow users to explore demographic and technical parameters that may influence the accuracy of local ancestry inference. However, because it uses the SELAM program to generate ancestry tract lengths (Corbett-Detig & Jones, 2016), it is also possible to implement natural selection during admixture (Figure S10). This will allow users to study the impacts of selection on local ancestry, ancestry junctions, and ancestry tract lengths. We predict that this will be a useful feature of *mixnmatch* for the many research groups studying selection after hybridization.

Simulated data from *mixnmatch* can be used to evaluate the accuracy of any ancestry inference program. However, it is designed to pair seamlessly with the *ancestryinfer* pipeline we describe here, which automates steps from read mapping to local ancestry inference. *ancestryinfer* has excellent accuracy under a broad range of biological conditions, and is fast and easy to use.

*ancestryinfer* is also intended to be an easy-to-use pipeline for local ancestry inference in real data. We demonstrate an application of the *ancestryinfer* pipeline to real data by using it to identify the locations of crossover events in *X. birchmanni* x *X. malinche* F_2_ hybrids (Figure 3) and constructing a recombination map (Figure 4). Other possible uses of *ancestryinfer* include generating ancestry probabilities for QTL mapping or for studying genome evolution in natural hybrid populations, highlighting the versatile applications of the *mixnmatch* and *ancestryinfer* pipelines.

## Data accessibility

Data associated with this manuscript is available on Dryad (doi:XXXX; Schumer, Powell, & Corbett-Detig, 2019) and *mixnmatch* and *ancestryinfer* pipelines are available on github (https://github.com/Schumerlab/mixnmatch; https://github.com/Schumerlab/ancestryinfer) and dockerhub (https://hub.docker.com/repository/docker/schumer/mixnmatch-ancestryinfer-image)

## Acknowledgements

We thank Peter Andolfatto, Andrés Bendesky, Quinn Langdon, Ben Moran, David Reich, Alisa Sedghifar, and members of the Schumer and Corbett-Detig labs for helpful discussions and feedback on this work. Stanford University and the Stanford Research Computing Center provided computational resources and support for this project. This work was supported by a Hanna H. Gray fellowship and NIH 1R35GM133774 grant to MS and by NIH 1R35GM128932 and an Alfred P. Sloan Fellowship to RC-D.

