## Supporting Information for "Versatile simulations of admixture and accurate local ancestry inference with *mixnmatch* and *ancestryinfer*"

#### Table of Contents

**Supporting Information 1.** Alternative hybrid simulation options in *mixnmatch* pipeline
**Supporting Information 2.** Information about configurable parameters in *ancestrinfer*
**Supporting Information 3.** Impact of reference bias on accuracy
**Supporting Information 4.** Complete description of *mixnmatch* simulations to evaluate accuracy
**Supporting Information 5.** Simulating RAD data and evaluating its impacts on accuracy
**Supporting Information 6.** Example of *mixnmatch* simulations with selection
**Supporting Information 7.** Data generation and ancestry inference in $F_1$ and $F_2$ hybrids between *X. birchmanni* and *X. malinche*
**Supporting Information 8.** Evaluating correlations between crossover and population genetic maps in swordtail fish

**Supporting Information Figures**
**Supporting Information References**

**Appendix 1.** User manual for *mixnmatch*
**Appendix 2.** User manual for *ancestryinfer*
**Appendix 3.** Protocol used for library preparation
**Appendix 4.** Example configuration files for specific simulations

**Supporting Information 1. Alternative hybrid simulation options in *mixnmatch* pipeline**

In the main text we describe our approach of simulating parental haplotypes and reference genomes using *macs* and *seq-gen.* We recommend using this approach to better model demographic history, background linkage disequilibrium, and shared and private polymorphisms in the parental species. However, because users may want to use the reference genomes and ancestry informative sites that they plan to use in downstream experiments, we provide an option for this in the *mixnmatch* pipeline (Figure S1).

With this option, users can provide two reference genomes and *mixnmatch* will identify candidate ancestry informative sites by tabulating all differences between them. These genomes must be in the same coordinate space for hybrid chromosome generation to occur properly. Because the provided reference genomes are only one sample from each parental population, we next simulate parental haplotypes by introducing variation into these sequences. First, *mixnmatch* uses the user-specified rate of shared polymorphism to convert a subset of differences between the reference sequences to shared polymorphisms. For guidance on determining the appropriate value for the rate of shared polymorphism parameter see Wakeley & Hey (1997). To be conservative, the frequency of shared polymorphisms is set at 0.5 in both parental species by default.

*mixnmatch* then calculates the expected number of polymorphic sites genome-wide for each parental species minus the re-assigned shared polymorphisms, and randomly selects the locations of these private polymorphisms from a uniform distribution. For each private polymorphism, we draw a frequency from the neutral site frequency spectrum. Finally, *mixnmatch* generates parental haplotypes based on the reference sequence for the appropriate species. In each individual at each polymorphic site the reference allele can be "mutated" to the alternative allele. Whether the alternative allele is introduced is determined with random binomial function, where the probability is set to the previously determined frequency of the non-reference allele.

We caution that although this approach incorporates observed ancestry informative sites, it assumes that both shared and private polymorphisms are distributed uniformly along the genome, ignores the effects of linkage within the source populations between such sites, and makes assumptions about the demographic history of the parental species (because it relies on the neutral site frequency spectrum). Thus, using this approach may result in an underestimation of the expected error rate.

**Supporting Information 2. Information about configurable parameters in *ancestrinfer***

Parameters for *ancestryinfer* are specified in a text-editable configuration file. Example configuration files can be found in Appendix 4 and more details on individual parameters can be found in the user manual (Appendix 2). Here, we highlight several important parameters that are user defined.

The HMM implemented through *ancestryinfer* (AncestryHMM, Corbett-Detig & Nielsen, 2017) can accept priors for mixture proportions and the time of initial mixture, or can attempt to infer these values from the data. If users wish to specify priors for these values, this can be done in the *ancestryinfer* configuration file. A recombination map can also be provided by the user; if a recombination map is not provided *ancestryinfer* will assume a uniform recombination prior.

An important step for local ancestry inference is the identification of ancestry informative sites to use in the HMM. If the two provided parental genomes are in the same coordinate space, users can direct *ancestryinfer* to identify ancestry informative sites between them. If users provide non-colinear genomes, a file containing the coordinates of ancestry informative sites must be provided in the configuration file. In this case *ancestryinfer* will use the coordinate space of the first parental genome provided (`"genome1"`) as the coordinate space for the HMM step. Thus, if there are significant differences in the quality of the two parental species' assemblies, we advise specifying the better assembly as genome 1.

Users can optionally specify a file that contains parental allele frequencies at ancestry informative sites. If no parental allele frequency counts are provided, *ancestryinfer* will assume that the sites that differ between the two genomes are fixed.

**Supporting Information 3. Impact of reference bias on accuracy**

One feature of *ancestryinfer* that is unusual is that it makes use of two reference genomes in an attempt to reduce artifacts of mapping biases. Briefly, reads from a hybrid individual are mapped to both species' reference genomes with *bwa* and mpileups are generated with *samtools*

(Li, 2011; Li & Durbin, 2009). Reads that do not map uniquely to both genomes are identified and removed from the bam file prior to mpileup generation. A similar approach is used in the multiplexed shotgun genotyping approach of Andolfatto et al. (2011) and recent approaches for quantifying allele specific expression (Zhang & Emerson, 2019).

We performed simulations to investigate whether this step reduced reference bias in the resulting count data. Because reference bias is predicted to be more problematic when the number of variant sites is high, we performed these analyses using simulations of species with 2% divergence and a 1% per-basepair polymorphism rate, similar to what is observed in some *Drosophila* species. We simulated 20 $F_1$ hybrids, which should generate equal counts mapping both species' reference genomes. We asked about the ratio of parent 1 to parent 2 reads using our approach versus using one or the other reference genome.

We found subtle but significant reference bias when mapping to either parental genome compared to the dual mapping approach described above. While 3-8% more reads mapped to the species 1 and species 2 genomes without the reference bias correction, this skew is reduced to 0.3-2% using our dual mapping approach, suggesting that it acts to mitigate the effects of reference bias. We note that when ancestral populations are closely related the effect of mapping bias on accuracy will likely be minor.

#### Supporting Information 4. Complete description of *mixnmatch* simulations to evaluate accuracy

To identify parameters that impact the accuracy of ancestry HMMs, we performed a number of simulations with *mixnmatch* and *ancestryinfer*. Although we focus here on evaluating performance based on the results of *ancestryinfer*, we note that the output files of *mixnmatch* (Table S1) can be used with any ancestry inference program.

As described in the main text and Table S2, our basic simulation scenario included the following parameters: we simulated 200 generations of admixture with a stable hybrid population size of 5,000 individuals, equal initial mixture proportions of the two parental species, a population mutation rate ($\theta$) in each of the parental species of 0.1%, and pairwise sequence divergence between the parental species of 0.5% ($D_{xy}$). We used a section of chromosome 1 from the swordtail fish *X. birchmanni* as the ancestral sequence to better simulate the variation in base composition and repeat content observed in real genomes. We also used the corresponding section of the *X. birchmanni* recombination map for chromosome 1 to specify recombination probabilities.

For computational speed, we simulated only 10 Mb of sequence (~1/3 the length of a typical swordtail chromosome), and 100 parental haplotypes of each species. Given the population mutation rate that we simulated (0.1%), this number of sampled parental haplotypes is expected to capture nearly all of the segregating sites within the parental species (Figure S2; (Watterson, 1975). We randomly sampled 20 haplotypes from each species to use in defining ancestry informative sites and required a frequency difference of 0.95 for a site to be treated as ancestry informative. We note that for less differentiated populations, such a stringent threshold could remove too many sites for accurate local ancestry inference so users should critically evaluate their choice of threshold. We generated 50 hybrid genomes for each simulation and simulated 0.5X coverage of these genomes. After each *mixnmatch* simulation we ran *ancestryinfer* and summarized accuracy at a posterior probability threshold of 0.9.

The results of these simulations indicated that accuracy under the baseline simulation conditions was expected to be excellent (Figure S4). This allowed us to explore in detail how changes in each of these parameters could influence the accuracy. For each simulation set we describe in the following sections, we used the parameter sets described above (listed in Table S2) but systematically varied values for one focal parameter. In the main text we describe the major results of these simulations; below we describe each simulation and its results in more detail.

*Accuracy as a function of the time since admixture*

In $F_1$ hybrids, individuals derive an entire chromosome from each of the parental species. As additional recombination events occur in subsequent generations, these ancestry tracts are broken down into smaller and smaller segments. At a certain point, ancestry tracts will become so small that they will be less effectively tagged by ancestry informative sites, resulting in a reduction in accuracy of local ancestry inference.

We evaluated how time since initial admixture influences accuracy by performing simulations of hybrid populations 500, 1000, 2500, and 5000 generations after initial admixture. The average ancestry tract length in these simulations ranged from 0.38 cM at generation 500 to

0.03 cM at generation 5000. We found that with increasing numbers of generations since initial hybridization, accuracy decreased (Figure 2). This effect was more pronounced in shorter ancestry tracts (Figure S5). Lower accuracy in shorter tracts is not surprising because these tracts will contain fewer ancestry informative sites. However, this observation does highlight the particular challenges associated with ancestry inference in ancient hybridization events. Moreover, these issues may be exaggerated in real data because regions of the genome with higher recombination rates tend to have fewer sites that are highly differentiated between species (as predicted by theory; Charlesworth, Charlesworth, & Morgan, 1995).

*Simulations with skewed admixture proportions*

Time since admixture is not the only factor that is expected to impact the lengths of ancestry tracts. When mixture proportions are unequal, theory predicts that this will change the mean ancestry tract length. Specifically, the length of ancestry tracts from the "minor" (i.e. less common) parental species will be shorter (Gravel, 2012). This suggests that mixture proportion may be another factor that influences accuracy in older hybrid populations.

To investigate this, we performed simulations of hybrid populations with older admixture (1,000 generations of admixture). We varied mixture proportions across simulations (70%, 80%, 90% and 95% parent species 1) and evaluated accuracy as a function of true ancestry state. While overall accuracy was similar across these simulations (99.2-99.5%), accuracy within ancestry tracts derived from the minor parent species was substantially lower as the skew in mixture proportion increased (Figure S11). This suggests that in hybrid populations with skewed admixture proportions error rates may differ as a function of ancestry state. Thus, it is important to be aware of these possible issues when analyzing ancestry variation along the genome.

*Accuracy as a function of divergence and genetic diversity in the parental species*

As the density of ancestry informative sites increases, accuracy of local ancestry inference is also expected to increase. Conversely, as the proportion of variant sites that are polymorphic in one or both parental species increases, accuracy is expected to decrease. The reason for this is two-fold: there will be a lower density of sites that have large frequency differences between species and more shared polymorphisms will be falsely assigned as ancestry informative sites.

Holding the rate of polymorphic sites constant, we simulated hybrids between species with increasing levels of divergence ($D_{xy}$ 0.25, 0.5, 0.75, and 1%). As discussed in the main text, accuracy dramatically increases with greater divergence between species, as does the spatial resolution of recombination events (Figure 2, Figure S3).

Next, we performed simulations holding the pairwise sequence divergence between species constant (at 0.5%) but increasing the level of genetic diversity within species. We accomplished this by modifying the population mutation rate ($\theta$) in simulations from 0.05% to 0.3%. As expected, we see that as within species diversity increases without a concurrent increase in divergent sites, the accuracy of local ancestry inference dramatically decreases (Figure S12). This decrease in accuracy is likely driven by a stark reduction in the number of ancestry informative sites used by *ancestryinfer*, falling from 0.3% of sites in the lowest polymorphism simulations to 0.06% in the highest polymorphism simulations (Figure S12).

*Simulating impacts of demographic history in the parental species*

The demographic history of each of the parental species will shape the genetic diversity within and divergence between these species at the time of hybridization. As a result, certain demographic scenarios might impact accuracy during local ancestry inference.

We first performed simulations as before but added changes in historical population size. Population expansions will result in an excess of rare variants in the parental species, while population bottlenecks can result in an excess of moderate or high frequency variants. We modeled an expansion in historical population size to 5 times the starting population size from 0.2*Ne* generations ago to the present day, where *Ne* is the effective population size. In a separate simulation, we modeled a population contraction to 0.2 times the starting population size from 0.2*Ne* generations ago to the present. Although we use these two scenarios as examples, we strongly recommend that users model demographic scenarios that match the inferred history of their focal populations.

Another process that is likely to have a major impact on the accuracy of local ancestry inference is historical migration between the two parental species. High levels of migration are predicted to reduce the number of ancestry informative sites, which will in turn lead to reduced accuracy. We simulated continuous, bidirectional migration between populations. Rates of migration ($m$) ranged from 0.005 to 0.05 across simulations and occurred from the time of the population split to the present.

We found that parental population contraction and expansion did not have a detectable impact on accuracy under the conditions simulated (Figure S13). However, this is likely to vary depending on the specific demographic scenario simulated. In contrast, we found that moderate to high levels of historical migration can significantly reduce accuracy in local ancestry inference in present-day hybrid populations (Figure S13). This reduction in accuracy is expected since high migration rates will reduce the number of sites that are differentiated between species. For example, in these simulations the number of ancestry informative sites at a differentiation threshold of 95% falls from ~0.25% per basepair with $m=0$ to ~0.15% per basepair with $m=0.05$.

Moreover, migration may result in more heterogeneity in the distribution of ancestry informative sites, impacting accuracy in low information regions. This highlights the importance of investigating expected accuracy under demographic scenarios that match the evolutionary history of their focal species.

We note that this effect could be compounded by selection against hybrids. Past research has shown that in many cases introgression rates are not uniform along the genome (Sankararaman, Mallick, Patterson, & Reich, 2016; Schumer et al., 2018). As a result, spatial heterogeneity in the number of ancestry informative sites along the genome (driven by historical hybridization) could bias inference in contemporary hybridization events.

*Simulating impacts of demographic history in the hybrid population*

After hybrid populations are formed, the demographic history of the hybrid population can impact the distribution of ancestry tract lengths. By default, if users do not provide information about the demographic history of the hybrid population, *mixnmatch* simulates a hybrid population with 5,000 diploid hybrid individuals, using the initial admixture proportions indicated in the configuration file. However, *mixnmatch* is capable of simulating arbitrarily complex demographic history within hybrid populations (Appendix 4). Users can specify this demographic history by providing a SELAM formatted demography file (Corbett-Detig & Jones, 2016) in the *mixnmatch* configuration file.

To demonstrate the application of this option in *mixnmatch*, we simulated a hybrid population with an ancient admixture event and a more recent admixture event. We simulated an initial hybrid population formed at 50-50 mixture proportions between the parental species 2,000 generations ago, with a burst of migration 200 generations ago from the same two parental species. This migration event replaced 50% of the hybrid population by adding 1,250 individuals from each parental species. We ran *ancestryinfer* with the option of allowing the HMM to infer the mixture time from the data.

As expected from the two admixture pulses simulated, ancestry tract lengths generated by this simulation show a bimodal distribution (Figure S14). Error rates from this simulation were also higher than in the simple admixture history evaluated in our base simulations (99±0.7%) but similar to those observed in simulations with older admixture (Figure 2). The *mixnmatch* and *ancestryinfer* configuration file and associated demography files used in this simulation are available in Appendix 4.

*Simulating impacts of drift in the reference panel*

To identify ancestry informative sites in the parental species and estimate their frequency a parental reference panel is required. Ideally this reference panel would be derived from the parental populations involved in the hybridization event, but this is not always possible. In many cases the source population itself could be admixed from ongoing hybridization, making it more practical to sample a non-admixed population as a reference panel. In other cases, the appropriate parental population may no longer be available, as is the case in the admixture events between humans and Neanderthals and Denisovans (Sankararaman et al., 2016).

We simulated reference parental populations with varying levels of divergence to the source populations involved in the hybridization event. For these simulations we varied the time of divergence from 0.1 to 0.75 (in units of *4Ne* generations). As in the base simulation, the total divergence time simulated between the parental species in *4Ne* generations was 2. An example configuration file for simulations of reference panel drift is available in Appendix 4. For simplicity, we simulated the same level of drift in both parental reference panels. We note, however, that *mixnmatch* accommodates simulations of drift in either parental population. Asymmetry in divergence to the reference panels could generate a scenario where ancestry calls for one parental species are less accurate.

We find that with low levels of drift between reference and source populations ancestry can be inferred with high accuracy in hybrids (Figure 2), but that error rates increase with increasing drift. This result is intuitive because sites that are ancestry informative in the reference panel may be found at different frequencies in the source parental populations, resulting in higher error rates.

Next, we explored whether this issue could be addressed in part by changing the required level of parental allele frequency differentiation. Because sites that are fixed or nearly fixed at the time of divergence between two populations of the same species are likely to remain so, increasing this threshold should improve accuracy when using drifted reference panels.

We simulated reference parental populations with high levels of drift from the admixing populations (0.75 in units of 4*Ne* generations) and tested accuracy at several differentiation thresholds (0.8, 0.9, 0.95 and 1). We find that accuracy improved significantly with more stringent allele frequency difference thresholds (Figure S15), suggesting this as a possible solution for researchers without access to the appropriate parental populations for generating reference panels.

*Simulating errors in the recombination map*

*ancestryinfer* and other local ancestry inference methods can take recombination priors that are then incorporated into transition probabilities in the HMM. Under ideal scenarios, incorporating recombination rate information is expected to increase accuracy in local ancestry inference. However, in practice, recombination maps are inferred with error, and these errors could outweigh any expected improvements in accuracy.

A common method for estimating local recombination rates is to use linkage disequilibrium (LD) based approaches from high coverage population samples (e.g. Auton & McVean, 2007; Chan, Jenkins, & Song, 2012). In a previous project, we estimated correlations between LD based recombination maps that were inferred from replicate simulations of 20 diploid individuals (Schumer et al., 2018). For these previous simulations, we modeled the same underlying recombination map using *macs* and a population mutation rate of 0.0012 per basepair (Schumer et al., 2018). The purpose of these simulations was to compare correlations between maps that were in fact identical to better understand the expected correlations between independently inferred LD-based maps. We found that although independently inferred LD-based maps were strongly correlated, substantial variation was introduced in the simulation and inference process (see Supporting Information of Schumer et al., 2018).

Here, we used these previous simulation results to generate an estimate of the degree of error expected in LD recombination maps. Although the expected error in LD recombination maps will vary based on focal species and sample size (Dapper & Payseur, 2018), because it is a commonly used approach for estimating recombination maps it serves as a useful example.

To generate an estimate of expected error, we calculated the standard deviation of differences in estimated recombination rate in the same SNP intervals from ten pairs of simulated maps. As expected, the mean difference in recombination rate across simulations was close to zero but the standard deviation was approximately $1^{-6}$ Morgans per interval.

We incorporated this estimate of the noise in map inference to investigate the impact of uncertainty in the recombination map on accuracy with *ancestyinfer*. We generated a focal dataset using *mixnmatch* and a prior recombination map (as used in base simulations), and simulated a population 2,500 generations post initial admixture, but otherwise used base parameters. We simulated an older admixture event because we reasoned that when ancestry tracts are smaller, the recombination prior may be more important in ancestry inference.

We first evaluated accuracy using the exact recombination map that was used in *mixnmatch* to simulate the data. Curious about how these results compared to a flat recombination prior, we re-ran the analysis, this time providing *ancestryinfer* with a uniform recombination map. Next, we wanted to evaluate the impacts of errors in the local recombination map on accuracy. Using the standard deviation inferred above, we added noise to the true recombination map by drawing from a normal distribution with a mean of zero and the observed standard deviation calculated above. Finally, because the simulations we used to estimate noise in map inference did not incorporate many possible sources of error (i.e. mapping, variant calling, etc), we repeated this procedure using a standard deviation two times greater.

We find that in the conditions simulated here, accuracy increases modestly but not significantly with a recombination prior in the limit of no map errors. However, adding realistic levels of noise expected from LD map inference can reduce accuracy (Figure S8). In practice, recombination map accuracy will depend on the method used to generate the map, resolution of the map, and quality of the data, among other factors. Users should thus evaluate what error rates might be expected in their study system and how this could influence accuracy in local ancestry inference.

*Simulating parameter misspecification*

In all of the simulations discussed above, we provided priors for admixture proportions and the number of generations since initial admixture to *ancestryinfer* that matched the true values simulated by *mixnmatch*. In practice, these parameters are estimated from data and are not known with certainty. To explore how sensitive *ancestryinfer* is to misspecification of input parameters, we performed a series of analyses.

Using our base simulation data, we misspecified the admixture time as 0.5 and 2 times the true admixture time in separate *ancestryinfer* runs respectively. We also evaluated the effects of misspecifying the admixture proportion by running *ancestryinfer* with admixture proportions of 0 and 1 (compared to the true mixture proportion of 0.5).

Surprisingly, we found that parameter misspecification did not have a detectable impact on the accuracy of *ancestryinfer* compared to *ancestryinfer* runs on the same data without misspecification (Figure S16). This suggests that *ancestryinfer* is not very sensitive to parameter misspecification. However, this may vary depending on the parameter misspecified or the degree of misspecification, so we recommend that researchers explore this question in their own simulations.

*Accuracy as a function of experimental design variables*

One of the major applications of *mixnmatch* is as a tool for researchers to use when making choices about experimental design for their projects. In this section we focus on choices researchers may make about sequencing effort for parental reference panels and for hybrid individuals.

One of the first decisions researchers will make when beginning a local ancestry inference project is how many individuals of each parental species to sequence to identify ancestry informative sites. The appropriate choice will vary as a function of genetic diversity within the parental species and divergence between them. To illustrate the importance of this choice, we performed simulations with base parameters but varied the number of parental haplotypes used to define ancestry informative sites (2, 5, 10, and 15 haplotypes of each species). When few parental haplotypes are used, many sites will be falsely identified as ancestry informative when they are in fact shared between the two species. We see this reflected in simulation results where accuracy is low when few parental haplotypes are sampled (Figure

S17). In practice, we expect diminishing returns beyond a certain sampling effort as most common polymorphisms will have been sampled (Figure S2).

Researchers will also have to decide how much data to collect for each hybrid individual. Because ancestry HMMs integrate information over consecutive ancestry informative sites, less coverage is required than for other applications such as variant calling. However, the required amount of coverage for high accuracy will depend on the age of the hybrid population, mixture proportion, and divergence between the parental species, among other parameters.

We performed simulations with base parameters, varying coverage from 0.05-1X per basepair genome-wide. Under these simulation conditions accuracy reached high levels even at per-base coverage of 0.25X (Figure 2). However, at lower coverage the resolution of the locations of recombination events was less precise, highlighting the utility of higher coverage in cases where researchers are interested in developing recombination maps (Figure S7).

*Simulating contamination*

Many popular library preparation protocols are performed in a plate based format (Andolfatto et al., 2011; Picelli et al., 2014). As a result, there is potential for cross-contamination between wells. Index hopping on some sequencing platforms can also generate reads that are mis-assigned to particular individuals (Valk, Vezzi, Ormestad, Dalén, & Guschanski, *bioRxiv*). These sources of contamination can introduce higher error rates, with the severity depending on the proportion of reads derived from the focal and contaminating individual.

If users specify a contamination rate in the configuration file, *mixnmatch* will add contamination as part of the read simulation step. This contamination rate will be used to determine the number of contaminant reads to add to the focal individual. Contaminant reads are generated from the full set of simulated parental haplotypes, and the proportion that match parent species 1 and parent species 2 is determined by the mixture proportion specified in the configuration file. A different set of contaminating reads is generated for each simulated hybrid.

We performed simulations of contamination guided by results from our empirical data. As part of our library preparation protocol (Appendix 3), we include twelve negative controls spread throughout the 96-well plate which are carried from DNA extraction through sequencing. This gives us an estimate of cross-well contamination in each plate (Figure S18). We used the average cross-well contamination rate observed in our empirical data (1.6%) with the base parameters used in other simulations.

We found that performance was relatively robust to these low levels of simulated contamination. However, at higher levels of simulated contamination (5-10%), accuracy decreased (Figure S19). Moreover, our simulations of uniform contamination rates do not capture the possibility that only certain individuals may have high levels of contamination which would introduce variance in accuracy across individuals.

**Supporting Information 5. Simulating RAD data and evaluating its impacts on accuracy**

Restriction site associated sequencing (or RADseq) has been a popular method for reduced representation sampling of the genome in ecology and evolutionary biology. However, because reduced representation sequencing will only sample a subset of ancestry informative sites in the genome and may introduce increased variation in the distance between these sites, it could have negative impacts on the accuracy of local ancestry inference. Moreover, with local ancestry inference methods that take advantage of count data, other features of RAD data could be problematic. Specifically, the goal of RADseq is often to generate pileups of reads at particular sites to facilitate variant calling, but variation in the depth of these pileups is expected due to polymorphisms that disrupt restriction enzyme target sites, as well as other sources of technical noise. This could generate variation in coverage in an allele specific manner that may interfere with HMMs that use count data, such as the one implemented in *ancestryinfer* (Corbett-Detig & Nielsen, 2017).

To evaluate this, we simulated RADseq data by generating reads only at sites following an EcoRI recognition sequence using a custom script (https://github.com/Schumerlab/Lab_shared_scripts). Otherwise, simulations followed the base parameter set used in other simulations (Supporting Information 4).

We found that accuracy with simulated RADseq data was substantially lower than with simulated low-coverage whole genome data (Figure S9). Examining the input to the HMM, we saw that sampled markers were relatively sparse throughout the simulated sequence, but have high per-individual coverage. This suggests that a more frequent cutter that more closely approximates low-coverage whole genome sequencing may have better performance, but that researchers should be cautious in using RAD data for local ancestry inference.

**Supporting Information 6. Example of *mixnmatch* simulations with selection**

Although we focus on simulating hybrid populations without selection in the main text, *mixnmatch* is designed to easily implement selection after hybridization. To do so, users simply need to specify a SELAM-formatted selection file in the *mixnmatch* configuration file (Corbett-Detig & Jones, 2016).

To demonstrate this, we performed simulations as described previously except that we simulated selection against parent 2 ancestry in the middle of the simulated sequence (0.1 cM position of the 0.2 cM simulated sequence). We simulated selection at $s$=0.25 on individuals homozygous for the parent 2 allele.

As expected, by generation 200 parent 2 ancestry in this region has been driven to low frequency in the hybrid population (Figure S10), and leaves a broad footprint of low parent 2 ancestry in the surrounding region. The configuration file from this example *mixnmatch* is included in Appendix 4. We expect that the option to implement selection will be a widely used feature of *mixnmatch* give recent interest in the impacts of selection on local ancestry patterns in hybrids.

**Supporting Information 7. Data generation and ancestry inference in $F_1$ and $F_2$ hybrids between *X. birchmanni* and *X. malinche***

To evaluate the performance of *ancestryinfer* on a known cross, we generated $F_1$ and $F_2$ hybrids between swordtail fish species *Xiphophorus birchmanni* and *X. malinche*. Pregnant wild caught *X. malinche* females from the Chicayotla population were allowed to give birth in captivity at Texas A&M University (IACUC protocol #2016-0190). Their offspring were raised and individuals were separated by sex as soon as males and females could be differentiated phenotypically. Virgin females from these lab-raised fish were combined with *X. birchmanni* males (collected from the Coacuilco *X. birchmanni* population) in 2000 liter tanks to produce $F_1$ offspring. After $F_1$ fry were born, adults were removed and fry were allowed to mature. The resulting $F_1$ fry from these tanks were distributed among two large mesocosm tanks to mature and mate. $F_2$ offspring from these $F_1$ tanks were split among four mesocosm tanks within their first two weeks of life and allowed to mature. Fin clips were taken from adult $F_1$ and $F_2$ fish.

We extracted DNA from fin clips of $F_1$ and $F_2$ artificial hybrids using the Agencourt DNAdvance kit following manufacturer's instruction. We generated low-coverage whole genome data for $F_1$ and $F_2$ hybrids using a Tn5 transposase based protocol (Appendix 3). We multiplexed individually barcoded libraries for all individuals and sequenced this pooled library on an Illumina HiSeq 2500 to collect paired-end 150 basepair reads. Reads were parsed by individual barcode using a custom script (https://github.com/Schumerlab/Lab_shared_scripts).

After parsing, these reads were used as input in the *ancestryinfer* pipeline. The reference genomes used with *ancestryinfer* were chromosome level *de novo* assemblies for *X. birchmanni* and *X. malinche*. The ancestry informative sites were identified in previous work (Schumer et al., 2018). We specified 2 generations as the prior for the time since initial admixture, a uniform recombination rate of 0.002 cM per kb, and the proportion of the genome from parent species 1 (in this case, *X. birchmanni*) as 0.5.

After running *ancestryinfer* we converted posterior probabilities for each ancestry state to hard-calls for a given ancestry using a posterior probability threshold of 0.9 and summarized admixture proportions and genome-wide heterozygosity at ancestry informative sites using a custom script (https://github.com/Schumerlab/Lab_shared_scripts/blob/master/parsetsv_ancestry_v2.pl).

For comparison, we simulated second generation hybrids with parameters matching those expected for hybrids formed between *X. birchmanni* and *X. malinche*. Specifically, we set the divergence time such that pairwise sequence divergence between the two species was 0.5% per basepair, simulated $\theta$ of 0.1% per basepair, and a bottleneck reducing one population to 0.25 its original size beginning 4*Ne* generations ago . This bottleneck roughly corresponds to a previously detected reduction in population size in *X. malinche* which has reduced present-day per basepair heterozygosity to 0.03% (Schumer et al., 2018). To match our real data, we simulated 0.2X coverage and 50-50 mixture proportions of the two parental species.

Results of analyses of $F_1$ and $F_2$ hybrids between *X. birchmanni* and *X. malinche* are summarized in Figure 3. We find extremely low rates of inferred ancestry homozygosity in $F_1$ hybrids (on average 0.003, 95% confidence intervals: 0.001 - 0.008). Importantly, the simulations of early generation hybrids with parameters chosen to mimic these artificial hybrids result in a similar estimated error rate within heterozygous tracts (on average 0.002, 95% confidence intervals: 0-0.006), with an average overall error rate of 0.002. Importantly, this comparison suggests that *mixnmatch* is generating simulated data with similar properties as real data.

**Supporting Information 8. Evaluating correlations between crossover and population genetic maps in swordtail fish**

Ancestry transitions observed in $F_2$ hybrid individuals reflect recombination events that occurred in their $F_1$ parents (Figure 3). Given our low estimated error rate, we predict that only a small subset of observed ancestry transitions are errors (e.g. Figure S20). Thus, we can use observed ancestry transitions in $F_2$ hybrids to build a genetic map.

We defined ancestry transitions as any switch in ancestry state between the three possible states in $F_2$ hybrids: homozygous parent 1 (*birchmanni*), heterozygous, or homozygous parent 2 (*malinche*). We defined the recombination breakpoint interval as the distance between the last high confidence (posterior probability >0.9) ancestry call for the previous ancestry state and the first high confidence ancestry call for the new ancestry state. Because we sequenced $F_2$ hybrids at relatively low coverage (0.2X), these intervals were quite large (lower and upper 5% quantile of interval size genome-wide: 23 kb - 667 kb). In addition to this uncertainty in the precise locations of recombination events, we also captured only 4,038 crossovers on 24 chromosomes in the 139 $F_2$ hybrids we sampled. As a result, we chose to summarize hybrid recombination rates in 5 Mb windows. Briefly, we summarized the total number of recombination events in the focal window as a proportion of the total events genome-wide in $F_2$ hybrids to generate a relative rate for each 5 Mb window.

We previously generated a LD based mapped for *X. birchmanni* using the program LDhelmet (Chan et al., 2012; Schumer et al., 2018). LDhelmet estimates the population recombination rate (ρ), which is the recombination rate scaled by 4*Ne*, and reports ρ/bp in SNP intervals along the genome. To convert these estimates to relative rates that we could compare to the $F_2$ map described above, we calculated a weighted average of rates between SNPs intervals and generated an overall estimate of the ρ/bp rate for each 5 Mb window. We compared the correlation between rate estimates across the two maps using a linear model in R. As expected, the crossover based map and the LD map were strongly correlated (R=0.82). The results of this comparison are shown in Figure 4.

Based on past work, we expect the true recombination maps (though not necessarily the inferred maps) to be nearly identical in *X. birchmanni, X. malinche*, and hybrids between the two species (Baker et al., 2017; Schumer et al., 2018). While the LD map reflects historical recombination events (and is influenced by a population's demographic history), the $F_2$ map only contains recombination events from the previous generation, making it much lower resolution. To understand if the observed correlation between the two maps is consistent with differences in resolution under the assumption that the underlying maps are in fact identical, we took a simple simulation-based approach.

For these simulations, we treated the rate in 5 Mb windows estimated in the LDhelmet map as the "true" rate. For each chromosome, we calculated the relative rate for each 5 Mb window in the LD map and the number of observed recombination events on that chromosome in $F_2$ hybrids. Next, we used the random multinomial function in R to draw recombination events for each window from the total number of events observed on that chromosome, using the relative rate from the LDhelmet map as the vector of probabilities. After performing this step for all chromosomes we recorded the correlation between the LD map and the simulated $F_2$ map. We repeated this procedure 500 times to obtain a distribution of expected correlations.

We find that the observed correlation between the LD and $F_2$ map falls within the range of what is expected from our simulations (Figure S21). We note that these simplistic simulations ignore sources of variation introduced by LD and $F_2$ map inference. However, this makes our simulations conservative as these sources of error would be expected to further reduce correlations between the two maps.

#### Supporting Information Tables

**Table S1.** Files produced by *mixnmatch* simulations for downstream use in local ancestry inference with *ancestryinfer* or other ancestry HMMs. File names contain information about the parameter sets used. X is substituted with the generations since initial admixture simulated and Y is substituted with the proportion of the hybrid population initially derived from the parent 1 species. N is substituted with the id number of the hybrid simulated.

| File or folder name | Description | Related files |
|---|---|---|
| simulated_hybrids_readlist_gen***X***_prop_par1_***Y*** | file containing path to the reads for use with *ancestryinfer* | Not applicable |
| simulated_hybrids_reads_gen***X***_prop_par1_ ***Y*** | folder containing reads, bed files indicating true ancestry state, and fasta files with the true nucleotide sequence for each hybrid haplotype | indiv***N***_read1.fq.gz<br>indiv***N***_read2.fq.gz<br><br>indiv***N***.bed<br><br>indiv***N***.fa |
| simulated_AIMs_for_AncestryHMM | file containing the identified ancestry informative sites for this simulation | Not applicable |
| simulated_parental_counts_for_AncestryHMM | Simulated parental allele frequencies and recombination rates assuming a uniform recombination prior at ancestry informative sites. *mixnmatch* also generates a version of this file with an informed recombination prior. | simulated_parental_counts_for_<br>AncestryHMM_recombination_prior.txt |
| macs_simulated_parent1.fa<br>macs_simulated_parent2.fa | Simulated parent 1 and parent 2 reference genomes | Not applicable |

**Table S2.** Base parameter values used in simulations. To evaluate how performance depended on individual parameters, one parameter of this base set was systematically varied. See Supporting Information 4 for descriptions of individual simulations.

| Parameter | Value in base simulation |
|---|---|
| Generations since admixture | 200 |
| Proportion of the genome from parent species 1 | 0.5 |
| Per basepair coverage | 0.5X |
| Diploid hybrid population size | 5,000 |
| Population mutation rate per basepair | 0.001 |
| Pairwise sequence divergence between parental species | 0.005 |
| Ancestral sequence | *X. birchmanni* chromosome 1 (1-10 Mb) |
| Recombination map | *X. birchmanni* chromosome 1 (1-10 Mb) |
| Number of parental haplotypes used | Total: 100 per species<br>Ancestry informative site definition: 20 per species |

#### Supporting Information Figures

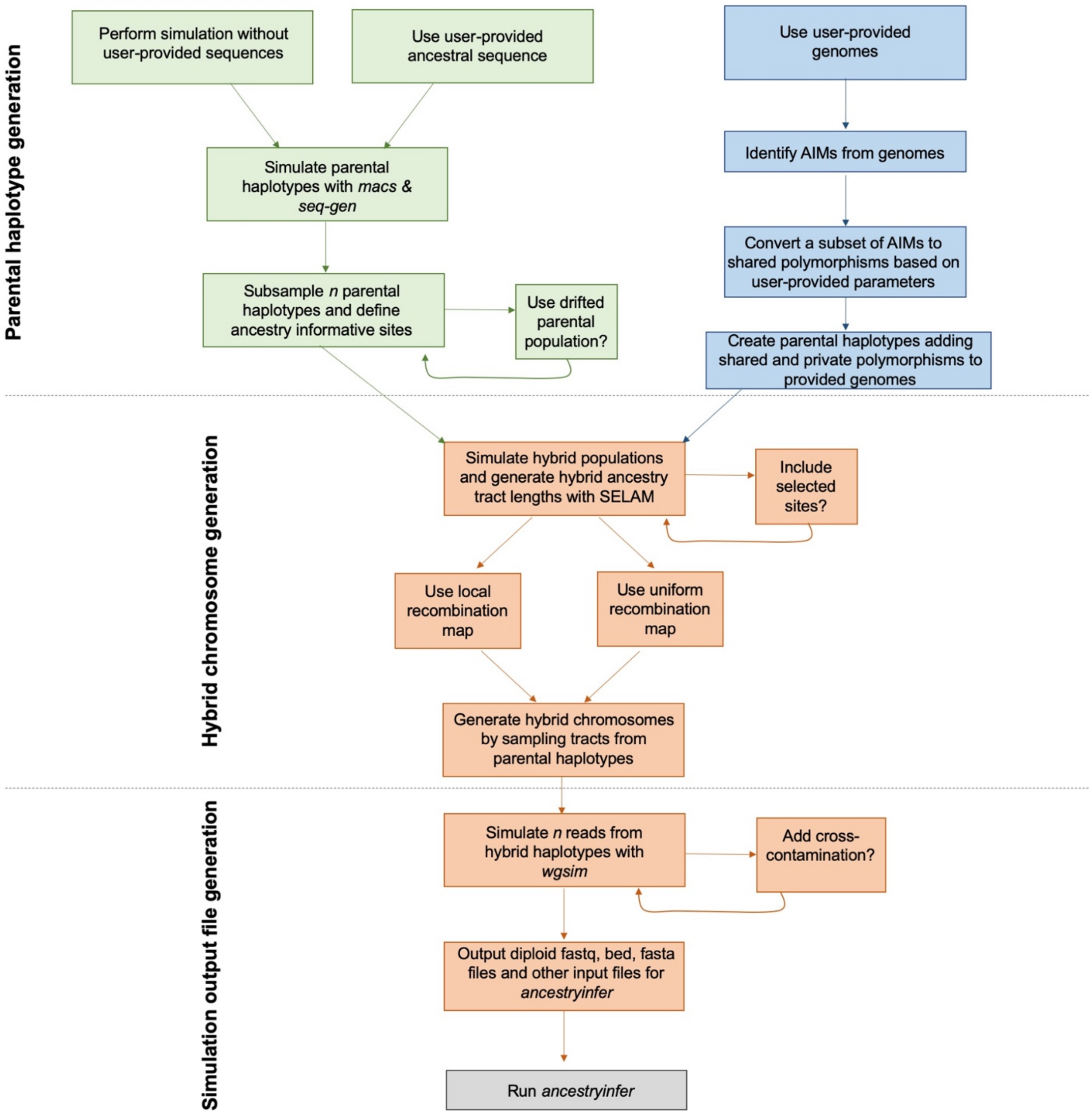


**Figure S1.** *mixnmatch* simulation flowchart indicating workflow and options available throughout the pipeline. All options can be specified in the text-editable configuration file (see Appendix 1, 4), Green boxes indicate the workflow using *macs* simulations of parental haplotypes (which can accommodate one user-provided genome as the ancestral sequence) and blue boxes indicate the workflow using two user-provided genomes without *macs*. Orange boxes indicate simulation steps downstream of parental haplotype generation that are shared by both modes of haplotype generation. AIMs is used in this figure as an abbreviation for ancestry informative markers. The final output of the *mixnmatch* pipeline is fastq and fasta files that can be input directly into *ancestryinfer* (Table S1).

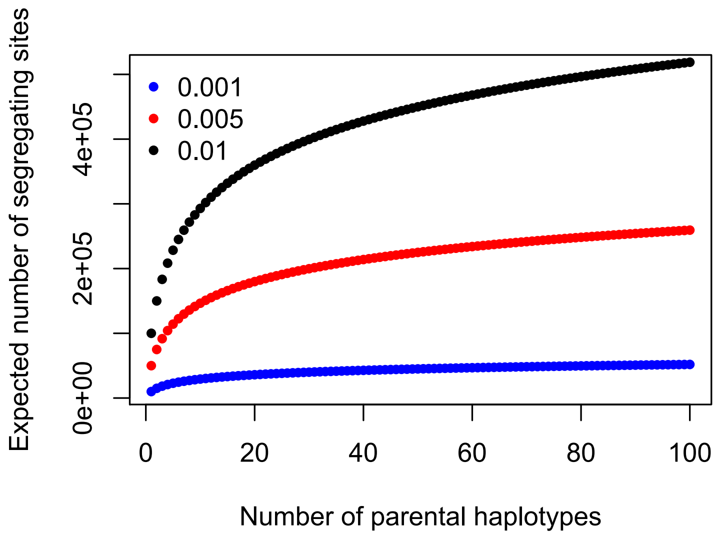


**Figure S2**. For the simulations described in the supplement and main text we generated 100 parental haplotypes from each species to use in simulations of hybrid genomes. This number of haplotypes is expected to capture much of SNP variation within the parental species, even in cases of high per basepair polymorphism. Shown here is the expected number of segregating sites detected in a 10 Mb segment at different per basepair polymorphism rates (0.001-0.01) as a function of the number of haplotypes sampled. Even in species with high rates of polymorphism, 100 haplotypes will capture much of the parental variation for the purposes of simulations.

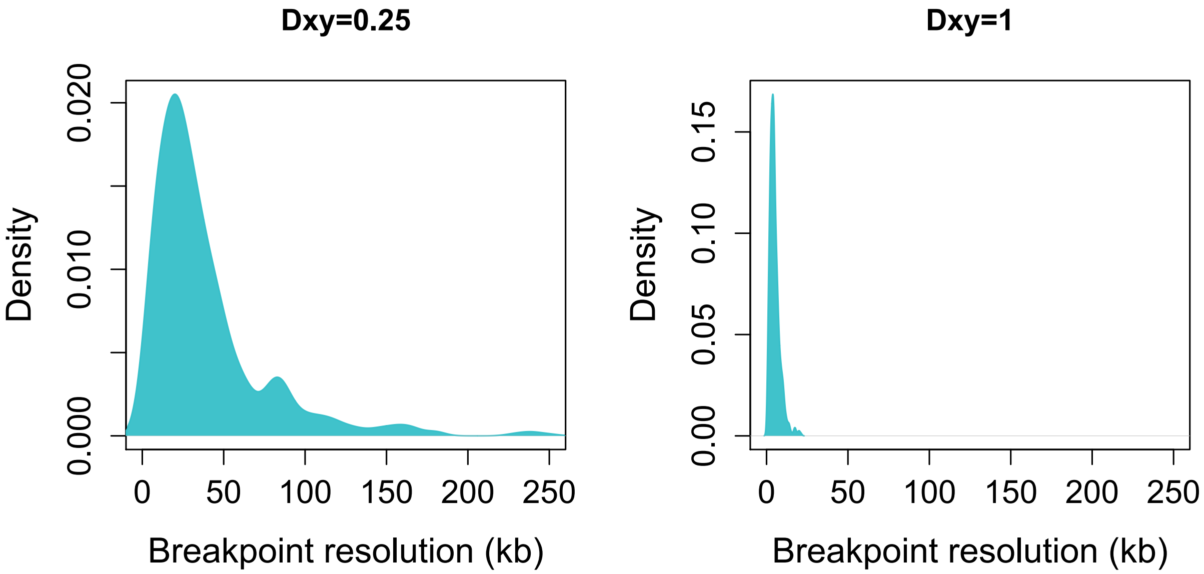


**Figure S3**. Another application of local ancestry data is using ancestry transitions to identify likely recombination events. For many applications the precise location of recombination events is important. The degree of sequence divergence between the parental species has a major impact on expected recombination breakpoint resolution (left - $D_{xy} = 0.25$, right - $D_{xy} = 1$). Recombination transitions are more precisely localized when ancestry informative sites are abundant.

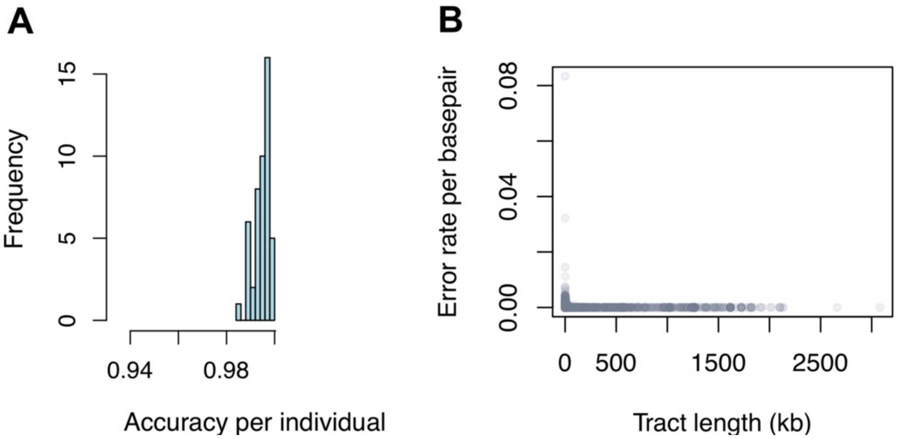


**Figure S4**. Example of output files summarizing accuracy that are generated by scripts provided with *mixnmatch*. The results shown here are from our base simulation conditions (Table S2; 200 generations since initial admixture, 50-50 mixture proportions, $D_{xy}$ = 0.5%, and per basepair polymorphism of 0.1%). A) The histogram shows the distribution of per-individual accuracy at ancestry informative sites genome-wide. B) A second plot produced by *mixnmatch* shows the relationship between tract length and accuracy, plotting ancestry tracts from all individuals.

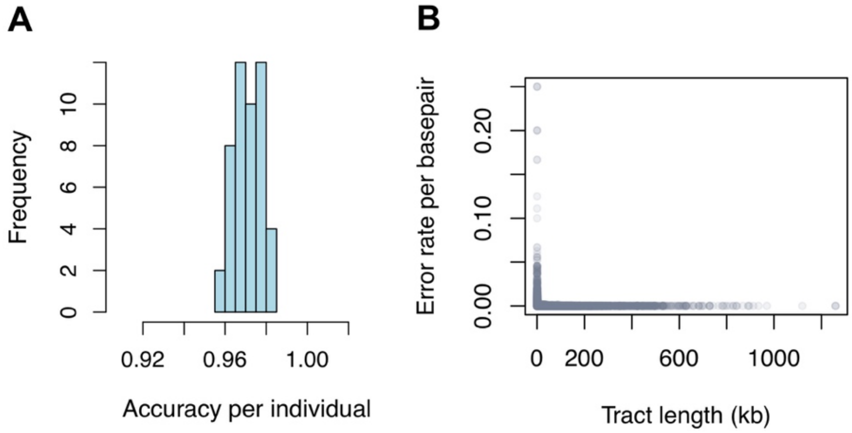


**Figure S5**. Per-individual accuracy (A) as well as accuracy in short ancestry tracts (B) decrease in simulations of older admixture events. The results shown here are from simulations of hybrid populations formed 5,000 generations ago (see also Figure 2) with other parameters set to values indicated in Table S2. Reduced individual level accuracy (A) in these simulations is in part attributable to high error rates in short ancestry tracts (B), presumably because too few ancestry informative sites exist in these tracts to accurately infer ancestry.

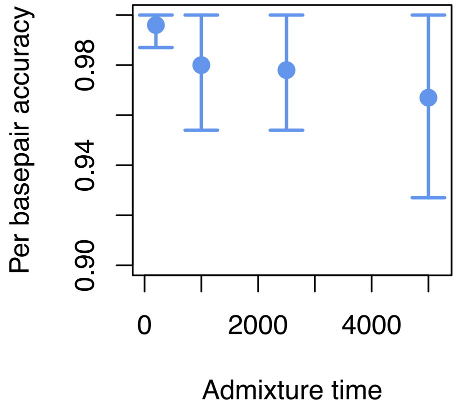


**Figure S6**. Accuracy versus admixture time in simulated populations that mix at 90:10 proportions. Both longer times since initial admixture and more skew in initial mixture proportion can result in shorter ancestry tracts. In these simulations we evaluated the impact of the time since initial admixture in populations that derived 90% of their genomes from one parental species and 10% from the other parental species. Compared to simulations with 50-50 mixture proportions, accuracy was lower in most simulations with skewed admixture proportions (Figure 2). Error bars show two standard deviations of individual-level accuracy in simulations. Note that asymmetry in error bars evident in some simulations is introduced by an upper limit of 100% accuracy.

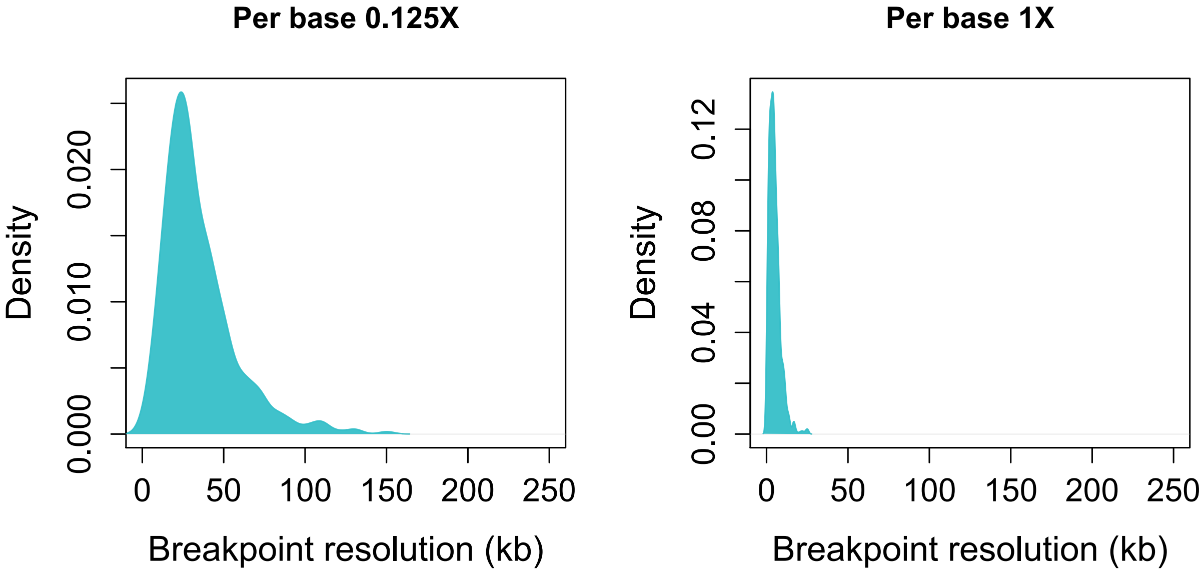


**Figure S7**. Recombination breakpoint resolution as a function of coverage. Although overall accuracy remains high in *mixnmatch* simulations with lower coverage (Figure 2), the locations of ancestry transitions are not as well resolved (left, average 0.125X coverage) as with higher coverage data (right, average 1X coverage).

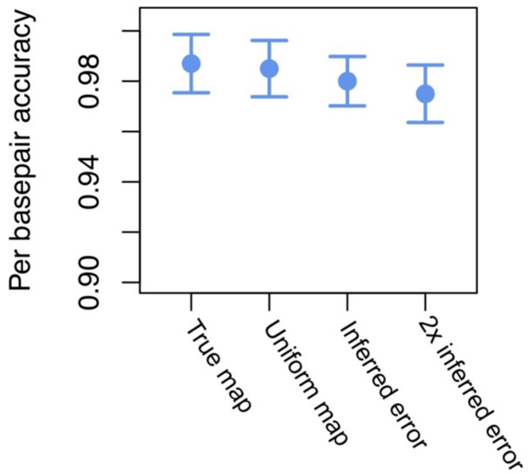


**Figure S8**. Results of simulations evaluating the impact of providing different recombination priors. We performed simulations with *mixnmatch* of admixture occurring 2,500 generations ago; a complete description of these simulations can be found in Supporting Information 4. We then evaluated accuracy when providing *ancestryinfer* with the true recombination map used in simulations, a uniform recombination map, and two maps with varying degrees of error introduced. In the first simulation with error, we used an estimate of the error rate expected from LD map inference (see Supporting Information 4). In the second, we evaluated the impact on accuracy with an error rate two times greater than this estimate. Error bars show two standard deviations of individual-level accuracy in simulations.

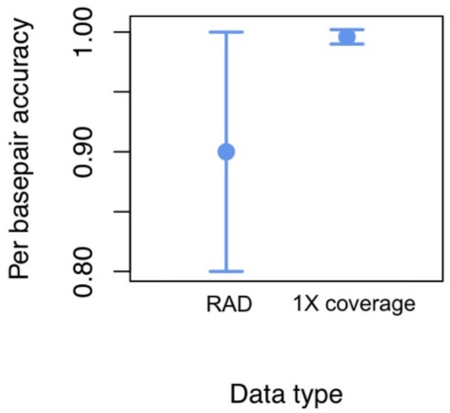


**Figure S9**. Comparison of accuracy of local ancestry inference using simulated RADseq and simulated low coverage whole genome data. Shown here is the results of local ancestry inference with data generated from the same set of simulated hybrid haplotypes. This analysis indicates that restriction digest associated data is expected to result in lower expected accuracy for local ancestry inference, likely because fewer ancestry informative markers are sampled (see Supporting Information 4). Error bars show two standard deviations of individual-level accuracy in simulations.

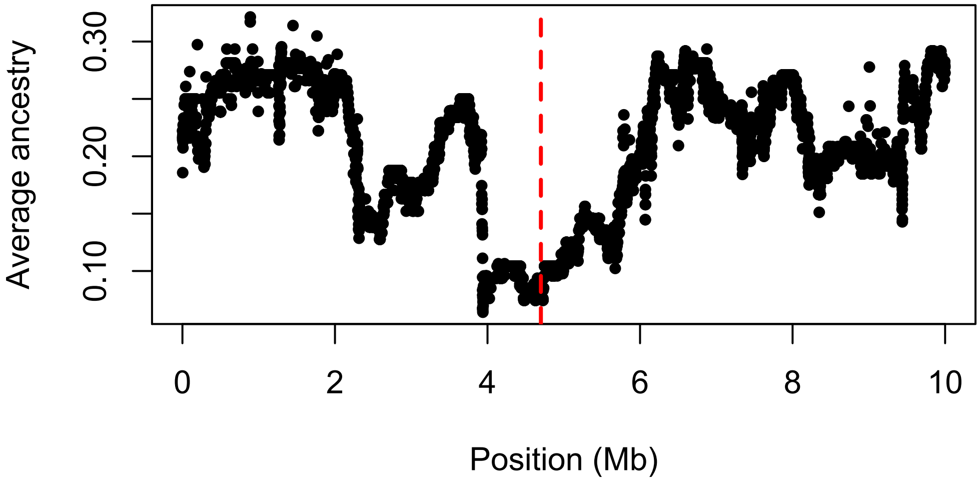


**Figure S10**. Example of selection on hybrids implemented with *mixnmatch*. Plot shows the average parent 2 ancestry across 50 individuals along the simulated 10 Mb region; the red line shows the location of the selected site. Simulated selection was against parent 2 ancestry in a population that formed at 50-50 mixture proportions 200 generations ago. Selection only occurred in individuals homozygous for parent 2 ancestry at the selected site.

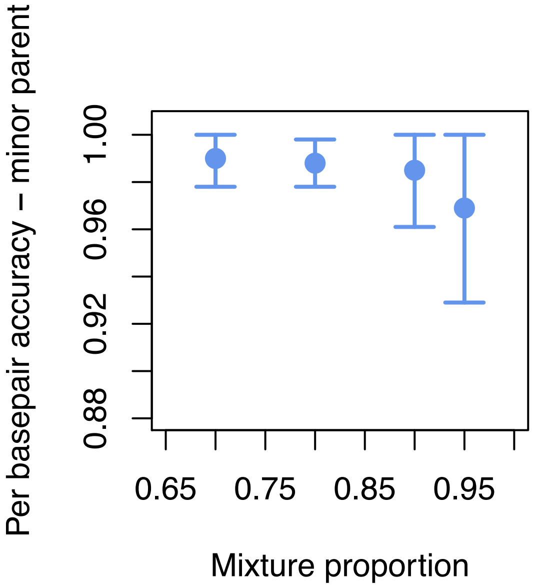


**Figure S11**. At skewed mixture proportions, ancestry tracts derived from the minor (i.e. less common) parental species are predicted to be shorter on average. This means that such tracts are tagged by fewer ancestry informative sites, which is predicted to result in lower accuracy in local ancestry inference. We observe this pattern in our simulations. Specifically, accuracy for the minor parent ancestry state decreases with increasing skew in mixture proportion, while accuracy for the major parent is unaffected. Error bars show two standard deviations of individual-level accuracy in simulations. Note that asymmetry in error bars evident in some simulations is introduced by an upper limit of 100% accuracy.

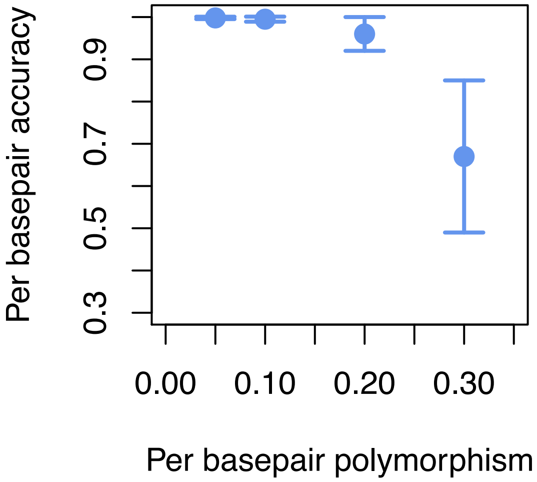


**Figure S12**. The accuracy of local ancestry inference is strongly dependent on the number of ancestry informative sites that distinguish the parental species (see Figure 2). This leads to the expectation that for a given rate of pairwise sequence divergence between species ($D_{xy}$), populations with higher rates of within species polymorphism will present more difficult cases for local ancestry inference. This pattern is indeed what we observe in simulations increasing rates of within species polymorphism (simulated $D_{xy}$ between species is held constant at 0.5%; Table S2). Error bars show two standard deviations of individual-level accuracy in simulations.

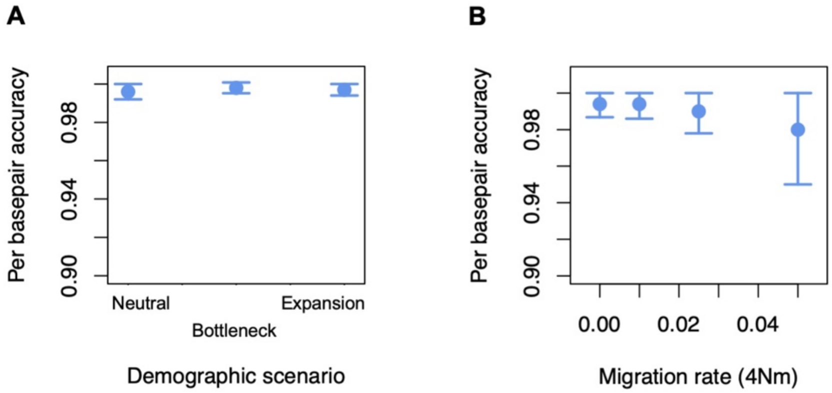


**Figure S13**. Example of simulations incorporating demographic events in the history of the parental species. While scenarios such as bottlenecks and population expansions did not have a major impact on accuracy under the parameters we simulated (A), high levels of migration between the parental species resulted in lower accuracy in local ancestry calling in present day hybrid populations (B). Error bars show two standard deviations of individual-level accuracy in simulations. Note that asymmetry in error bars evident in some simulations is introduced by an upper limit of 100% accuracy.

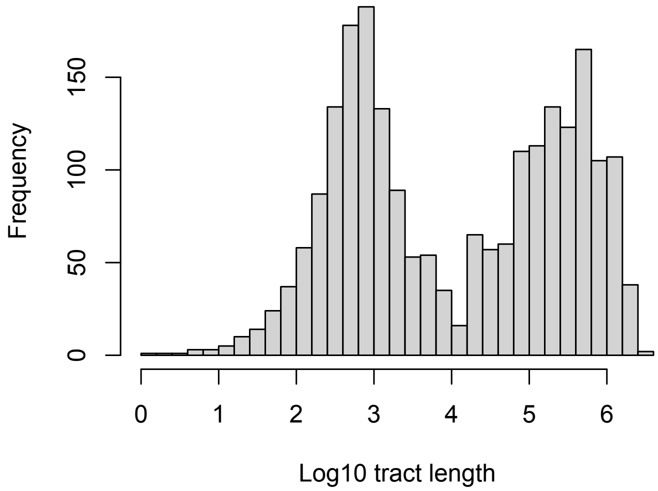


**Figure S14**. Ancestry tract length distribution from a *mixnmatch* simulation with two pulses of admixture. In this simulation we modeled two pulses of admixture, one 2,000 generations ago and a recent pulse 200 generations ago (see Supporting Information 4 for details). As expected, this generates a bimodal distribution of ancestry tracts, reflecting the two admixture pulses.

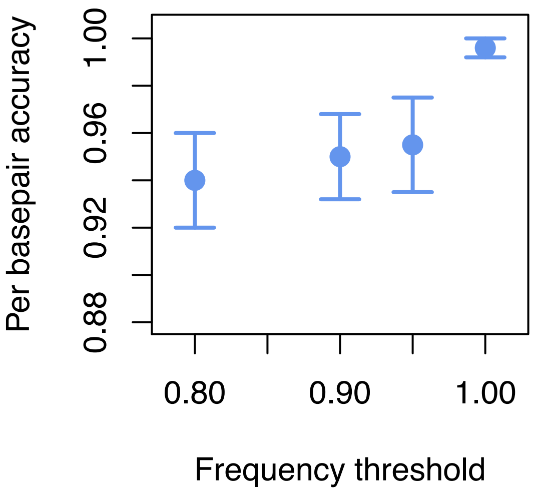


**Figure S15**. Genetic drift between the sampled parental populations and those that actually participated in the hybridization event is expected to reduce accuracy in local ancestry inference (Figure 2). Our simulations suggest that this can be mediated in part by imposing higher thresholds for allele frequency differences between parental populations. We simulated high levels drift between the reference and hybridizing parental populations (0.75 in units of *4Ne* generations, or ~35% of the branch separating the two species). For each simulation, we applied a different required threshold for allele frequency differences between the reference parental populations. Result suggest that high frequency thresholds can in part mediate issues introduced by using a drifted reference population. Error bars show two standard deviations of individual-level accuracy in simulations.

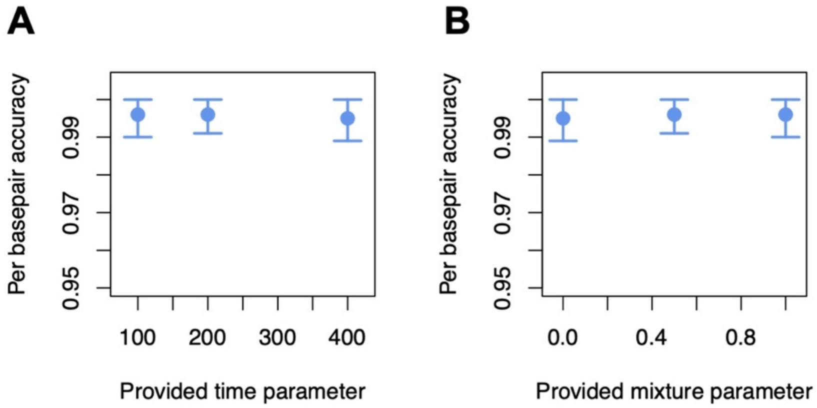


**Figure S16**. Users have the option of providing estimates for some parameters to *ancestryinfer*, but these parameters may be misspecified if users cannot accurately estimate their values. We simulated misspecification of admixture time (A, true value: 200) and admixture proportion (B, true value: 0.5). Surprisingly, we found that misspecification of these parameters did not have a detectable impact on accuracy, though this is likely to be sensitive to simulation conditions. Error bars show two standard deviations of individual-level accuracy in simulations.

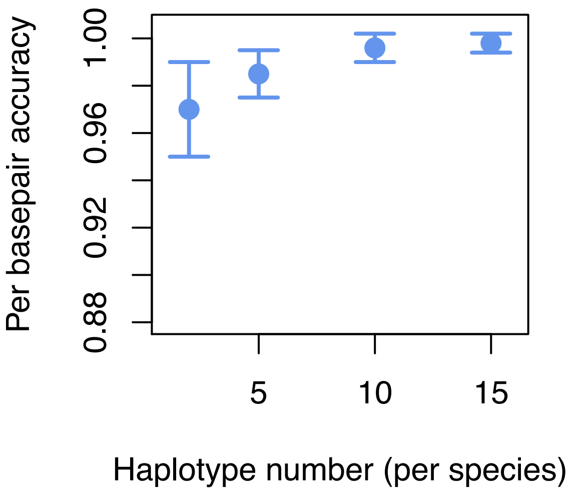


**Figure S17.** Accuracy as a function of the number of parental haplotypes used in defining ancestry informative sites. In our base simulations, we use 20 parental haplotypes from each parental species to define ancestry informative sites (Table S2). The accuracy of local ancestry inference is expected to decrease as the number of parental haplotypes sampled decreases, because with few haplotypes sites that are segregating in one or both of the parental species are more likely to be falsely assigned as ancestry informative. This prediction is borne out in simulations sampling 2-15 parental haplotypes per species. Error bars show two standard deviations of individual-level accuracy in simulations.

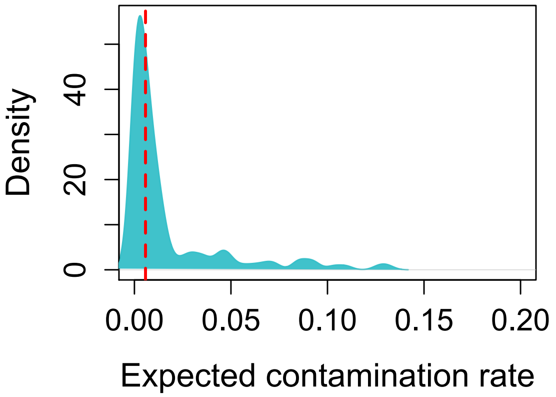


**Figure S18**. Empirical estimates of cross-well contamination rate based on 200 control samples over nine sequencing runs. Coverage in the control wells was normalized to average coverage of focal samples in the sequencing run. Negative controls were embedded from the DNA extraction through the sequencing stage, giving us estimates of cross well contamination at several sites across the plate. The dashed red line shows the median estimated contamination rate of 0.016.

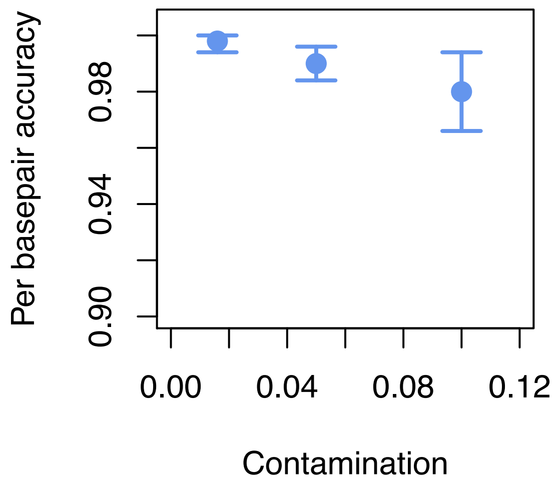


**Figure S19**. Accuracy as a function of simulated contamination rate using *mixnmatch*. While error rates are low at simulated levels of contamination close to what is observed in our empirical data (median 0.016), error rates increase with increasing contamination rates. Error bars show two standard deviations of individual-level accuracy in simulations.

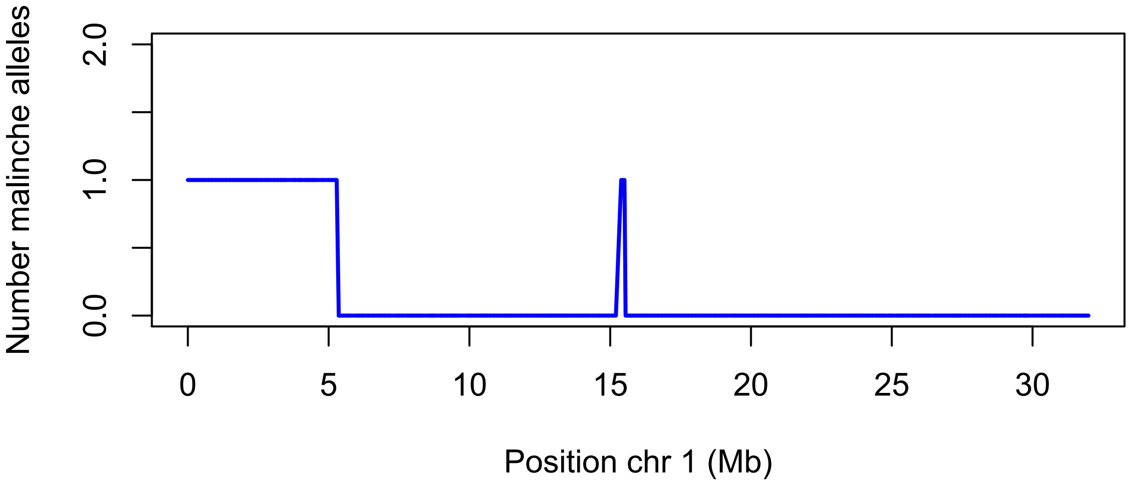


**Figure S20**. Errors in both simulated and empirical data occur at a low rates and can often be visually detected as short switches in ancestry state. Here we show a likely error detected in an $F_2$ hybrid individual on chromosome 1. Seven ancestry informative markers (out of ~34,000 chromosome wide) in the middle of the chromosome switch ancestry from homozygous *birchmanni* to heterozygous.

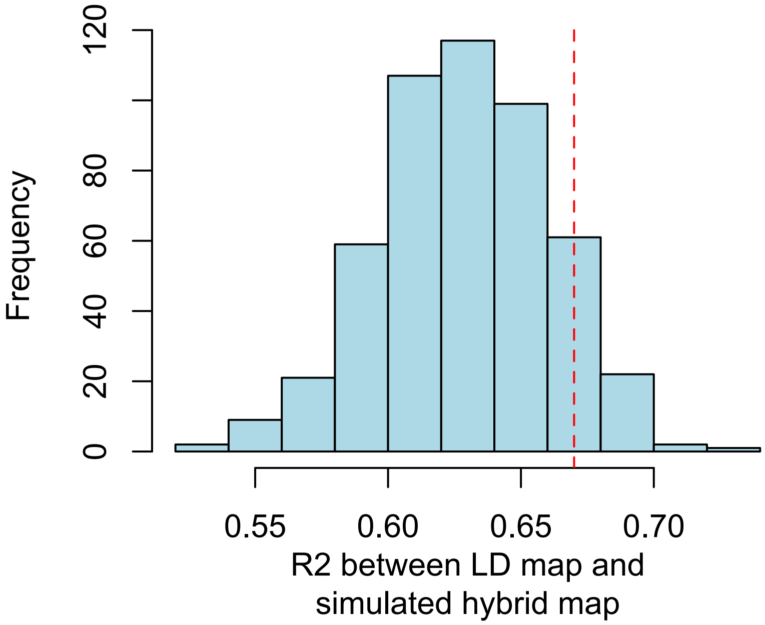


**Figure S21**. Results of simulations to determine the expected correlations between the LD and $F_2$ maps, assuming that the two maps are in fact identical. In these simulations we counted the number of observed ancestry transitions in $F_2$ hybrids. We then simulated new $F_2$ maps using the estimated rates from the LD map for *X. birchmanni* and summarized recombination rates in 5 Mb windows (see Supporting Information 8). We found that the observed level of correlation between the LD and $F_2$ maps (shown by the red dashed line) is consistent with expectations from these simulations. We note that although these simple simulations neglect important sources of variation they are likely conservative for the purposes of this comparison (see discussion in Supporting Information 8).

#### Supporting Information References


Andolfatto, P., Davison, D., Erezyilmaz, D., Hu, T. T., Mast, J., Sunayama-Morita, T., & Stern, D. L. (2011). Multiplexed shotgun genotyping for rapid and efficient genetic mapping. *Genome Research*, *21*(4), 610–617. doi: 10.1101/gr.115402.110

Auton, A., & McVean, G. (2007). Recombination rate estimation in the presence of hotspots. *Genome Research*, *17*(8), 1219–1227. doi: 10.1101/gr.6386707

Baker, Z., Schumer, M., Haba, Y., Bashkirova, L., Holland, C., Rosenthal, G. G., & Przeworski, M. (2017, June 6). Repeated losses of PRDM9-directed recombination despite the conservation of PRDM9 across vertebrates. doi: 10.7554/eLife.24133

Chan, A. H., Jenkins, P. A., & Song, Y. S. (2012). Genome-Wide Fine-Scale Recombination Rate Variation in Drosophila melanogaster. *PLOS Genetics*, *8*(12), e1003090. doi: 10.1371/journal.pgen.1003090

Charlesworth, D., Charlesworth, B., & Morgan, M. T. (1995). The pattern of neutral molecular variation under the background selection model. *Genetics*, *141*(4), 1619–1632.

Corbett-Detig, R., & Jones, M. (2016). SELAM: simulation of epistasis and local adaptation during admixture with mate choice. *Bioinformatics*, *32*(19), 3035–3037. doi: 10.1093/bioinformatics/btw365

Corbett-Detig, R., & Nielsen, R. (2017). A Hidden Markov Model Approach for Simultaneously Estimating Local Ancestry and Admixture Time Using Next Generation Sequence Data in Samples of Arbitrary Ploidy. *PLOS Genetics*, *13*(1), e1006529. doi: 10.1371/journal.pgen.1006529

Dapper, A. L., & Payseur, B. A. (2018). Effects of Demographic History on the Detection of Recombination Hotspots from Linkage Disequilibrium. *Molecular Biology and Evolution*, *35*(2), 335–353. doi: 10.1093/molbev/msx272

Gravel, S. (2012). Population Genetics Models of Local Ancestry. *Genetics*, *191*(2), 607. doi: 10.1534/genetics.112.139808

Li, H. (2011). A statistical framework for SNP calling, mutation discovery, association mapping and population genetical parameter estimation from sequencing data. *Bioinformatics*, *27*(21), 2987–2993. doi: 10.1093/bioinformatics/btr509

Li, H., & Durbin, R. (2009). Fast and accurate short read alignment with Burrows-Wheeler transform. *Bioinformatics*, *25*(14). doi: 10.1093/bioinformatics/btp324

Picelli, S., Björklund, Å. K., Reinius, B., Sagasser, S., Winberg, G., & Sandberg, R. (2014). Tn5 transposase and tagmentation procedures for massively scaled sequencing projects. *Genome Research*, *24*(12), 2033–2040. doi: 10.1101/gr.177881.114

Sankararaman, S., Mallick, S., Patterson, N., & Reich, D. (2016). The Combined Landscape of Denisovan and Neanderthal Ancestry in Present-Day Humans. *Current Biology*, *26*(9), 1241–1247. doi: 10.1016/j.cub.2016.03.037

Schumer, M., Xu, C., Powell, D. L., Durvasula, A., Skov, L., Holland, C., … Przeworski, M. (2018). Natural selection interacts with recombination to shape the evolution of hybrid genomes. *Science*, *360*(6389), 656. doi: 10.1126/science.aar3684

Valk, T. van der, Vezzi, F., Ormestad, M., Dalén, L., & Guschanski, K. (n.d.). Index hopping on the Illumina HiseqX platform and its consequences for ancient DNA studies. *Molecular Ecology Resources*, *0*(0). doi: 10.1111/1755-0998.13009

Wakeley, J., & Hey, J. (1997). Estimating Ancestral Population Parameters. *Genetics*, *145*(3), 847–855.

Watterson, G. A. (1975). On the number of segregating sites in genetical models without recombination. *Theoretical Population Biology*, *7*(2), 256–276.

Zhang, X., & Emerson, J. J. (2019). Inferring the genetic architecture of expression variation from replicated high throughput allele-specific expression experiments. *BioRxiv*, 699074. doi: 10.1101/699074

### Appendix 1: *mixnmatch* simulator user manual

#### Getting Started

##### Install

*Option 1 – install dependencies:*

```
git clone https://github.com/Schumerlab/mixnmatch.git
```

To install dependencies, follow instructions outlined in:

```
installation_instructions.txt
```

Test that the install and pipeline are working:

```
cd mixnmatch

perl simulate_admixed_genomes_v6.pl
hybrid_simulation_configuration_ancestral_seq_example_nonparallel.cfg
```

*Option 2 – load docker file for dependencies:*

With docker:

```
docker pull schumer/mixnmatch-ancestryinfer-image:mixnmatch-
ancestryinfer-docker

docker run -it mixnmatch-ancestryinfer-image bash
```

Test that the install and pipeline are working:

```
cd mixnmatch

perl simulate_admixed_genomes_v6.pl
hybrid_simulation_configuration_ancestral_seq_example_nonparallel.cfg
```

With singularity:

```
singularity pull docker://schumer/mixnmatch-ancestryinfer-
image:mixnmatch-ancestryinfer-docker

git clone https://github.com/Schumerlab/mixnmatch.git

singularity run mixnmatch-ancestryinfer-image_mixnmatch-ancestryinfer-
docker.sif bash
```

Test that the install and pipeline are working:

```
cd mixnmatch

perl simulate_admixed_genomes_v6.pl
hybrid_simulation_configuration_ancestral_seq_example_nonparallel.cfg
```

**Setting parameters in the configuration file**

There are several example configuration files available from github:

1) Example of a basic macs-based simulation using a user-provided ancestral sequence

`hybrid_simulation_configuration_ancestral_seq_example_parallel.cfg`

`hybrid_simulation_configuration_ancestral_seq_example_nonparallel.cfg`

2) Example of a macs-based simulation with drift between the hybridizing populations and the reference parental populations

`hybrid_simulation_configuration_sourcepopdrift_example_parallel.cfg`

3) Example of a simulation with user-provided reference genomes

`hybrid_simulation_configuration_usergenomes_example_parallel.cfg`

Parameter descriptions:
*Note: some parameters should only be set if you are using macs to simulate parental genomes and some should only be set if you are using your own genomes. See below sections for a list of each.*

| **Parameter** | **Description** | **Example** | **Include if** |
|---|---|---|---|
| `genome1=` | User provided fasta file for species 1 or ancestral sequence | `genome1=group1_par1.fa` | `use_ancestral=1`<br>or<br>`use_macs=0` |
| `genome2=` | User provided fasta file for species 2 | `genome2= group1_par2.fa` | `use_macs=0` |
| `use_ancestral=` | Treat genome1 as an ancestral sequence in | `use_ancestral=1` | Optional to include if `use_macs=1` |

| | | | |
|---|---|---|---|
| | simulations (options are 0 - no or 1 - yes) | | |
| mixture_prop_par1= | Expected proportion of the genome derived from parent species 1 | mixture_prop_par1=0.5 | Always required |
| rec_rate_Morgans_kb= | Recombination rate in Morgans per kb to simulate | rec_rate_Morgans_kb=0.00002 | Always required |
| num_indivs= | Number of individuals to simulate | num_indivs=50 | Always required |
| gens_since_admixture= | Generations since initial admixture to simulate | gens_since_admixture=50 | use_macs=0<br><br>or<br><br>SELAM_param_file= |
| chr_to_simulate= | Chromosome to use for simulations | chr_to_simulate=group1 | use_macs=0 |
| poly_perbp_par1= | Per-basepair polymorphism rate in parent species 1 | poly_perbp_par1=0.001 | use_macs=0 |
| poly_perbp_par2= | Per-basepair polymorphism rate in parent species 2 | poly_perbp_par2=0.001 | use_macs=0 |
| rate_shared_poly_at_aims= | Rate of shared polymorphisms between species | rate_shared_poly_at_aims=0.01 | use_macs=0 |
| read_type= | Type of reads to simulate (paired end - PE or single end - SE) | read_type=PE | Always required |

| | | | |
|---|---|---|---|
| `read_length=` | Length of reads to simulate | `read_length=100` | Always required |
| `per_bp_indels=` | INDEL rate per basepair to simulate in reads | `per_bp_indels=0.006` | Always required |
| `sequencing_error=` | Sequencing error rate per basepair to simulate | `sequencing_error=0.005` | Always required |
| `number_reads=` | Number of reads to simulate | `number_reads=100000` | Always required |
| `parental_drift=` | Simulate drift from the source parental populations (0 - no, 1 - yes). **Note: this must be paired with the appropriate macs command** | `parental_drift=0` | Required if the following parameters are defined: `macs_par1_aims_pop=` `macs_par2_aims_pop=` |
| `macs_par1_aims_pop=` | Parent 1 population to sample for AIMs generation | `macs_par1_aims_pop=3` | Required if `parental_drift=1` |
| `macs_par2_aims_pop=` | Parent 2 population to sample for AIMs generation | `macs_par2_aims_pop=4` | Required if `parental_drift=1` |
| `aim_freq_cutoff=` | Frequency difference required between parental populations to treat a site as ancestry informative | `aim_freq_cutoff=0.9` | Always required |
| `cross_contam=` | Contamination rate to simulate in | `cross_contam=0.02` | Optional |

| | | | |
|---|---|---|---|
| | hybrids. Contamination reads are drawn from the parental haplotypes at the observed mixture proportion. | | |
| `job_submit_cmd=` | Use slurm resource management system or run individuals sequentially | `job_submit_cmd=sbatch`<br><br>or<br><br>`job_submit_cmd=bash` | Always required. If sbatch is specified, users must provide a job submission header in `job_header=` |
| `use_macs=` | Indicate whether to use macs with seq-gen to simulate sequences (0 - no, 1 - yes) | `use_macs=1` | Always required |
| `use_map=` | Use a macs-formatted local recombination map in simulations of parental and hybrid populations and to set recombination priors (0- no, 1 - yes) | `use_map=1` | Always required |
| `SELAM_param_file=` | Provide a parameter file for SELAM describing the hybrid population history. If this is left | `SELAM_param_file=`selam_demography_params.txt | Only required if users wish to simulate a particular demographic history in hybrids |

| | | | |
|---|---|---|---|
| | blank the program assumes a neutral demographic history. | | |
| SELAM_selection_file= | Provide a SELAM formatted file indicating which site experience selection. See SELAM documentation. | SELAM_selection_file=selam_selection.txt<br><br>Example:<br><br>$cat selam_selection.txt<br>S A 0 0.1 1 1 0.9 | Optional |
| macs_params= | Command to be used for macs simulations of parental species demographic history. See ms/macs documentation for simulation options. | macs_params=200 10000000 -I 2 100 100 0 -t 0.001 -h 1e2 -r 0.001 -ej 2 2 1 -R recombination_map_for_macs.txt<br><br>macs_params=80 10000000 -I 4 20 20 20 20 0 -t 0.001 -h 1e2 -r 0.001 -ej 0.05 3 2 -ej 0.051 4 1 -ej 2 2 1 -R recombination_map_for_macs.txt | Required if use_macs=1 |
| program_path= | Path to the install location of the simulator program. If left blank, the program assumes a global install. | program_path=/home/bin | Optional |
| par1_for_aims= | Number of parent 1 haplotypes to use to define ancestry | par1_for_aims=20 | Required if use_macs=1 |

| | | | |
|---|---|---|---|
| | informative sites | | |
| par2_for_aims= | Number of parent 2 haplotypes to use to define ancestry informative sites | par2_for_aims=20 | Required if use_macs=1 |
| seq_params= | Provide base composition and transition transversion parameters for seq-gen sequence generation | | Optional if use_macs=1 |
| job_header= | Provide cluster-specific job submission parameters to be used to submit simulation jobs | job_header=#!/bin/sh #SBATCH --ntasks=1 #SBATCH --cpus-per-task=1 --mem=64000 #SBATCH --time=02:00:00 | Always required |
| num_indiv_per_job= | Set how much parallelization to perform by setting the number of individuals to run per submitted job. To run a different job for each individual set this parameter to 1 | num_indiv_per_job=2 | Always required |

We also provide *macs* and SELAM documentation in the git repository for *mixnmatch* for convenience.

**Examples**

Several example files are available with the git repository including example configuration files

**Running the pipeline**

After setting the parameters in the configuration file and loading required dependencies, simply run:

```
perl mixnmatch/ simulate_admixed_genomes_v6.pl
hybrid_simulation_configuration.cfg
```

where path is the path to your simulator install

#### Appendix 2 *ancestryinfer* pipeline

Output of *mixnmatch* simulations or your own data can be input into the *ancestryinfer* pipeline to run local ancestry inference following (Corbett-Detig & Nielsen, 2017).

##### Install

*Option 1 – install dependencies:*

```
git clone https://github.com/Schumerlab/ancestryinfer.git
```

To install dependencies, follow instructions outlined in:

```
installation_instructions.txt
```

Test that the install and pipeline are working:

```
cd ancestryinfer

perl Ancestry_HMM_parallel_v5.pl
hmm_configuration_file_nonparallel.cfg
```

*Option 2 – load docker file for dependencies:*

With docker:

```
docker pull schumer/mixnmatch-ancestryinfer-image:mixnmatch-
ancestryinfer-docker

docker run -it mixnmatch-ancestryinfer-image bash
```

With singularity:

```
singularity pull docker://schumer/mixnmatch-ancestryinfer-
image:mixnmatch-ancestryinfer-docker

git clone https://github.com/Schumerlab/ancestryinfer.git

singularity run mixnmatch-ancestryinfer-image_mixnmatch-ancestryinfer-
docker.sif bash
```

Test that the install and pipeline are working:

```
cd ancestryinfer
```

```
perl Ancestry_HMM_parallel_v5.pl
hmm_configuration_file_nonparallel.cfg
```

**Setting parameters in the configuration file**

There is are example configuration files available on github:

`hmm_configuration_file_parallel.cfg`

`hmm_configuration_file_nonparallel.cfg`

Parameter descriptions:

| Parameter | Description | Example | Include if |
|---|---|---|---|
| `genome1=` | User provided fasta file for species 1 | `genome1=xiphophorus_birchmanni_10x_12Sep2018_yDAA6.fasta` | Always |
| `genome2=` | User provided fasta file for species 2 | `genome2=Xmalinche_dovetail_assembly.fa` | Always |
| `read_type=` | Indicated whether data is paired end or single end | `read_type=PE` | Always |
| `read_list=` | Provide list (including full paths) to the reads to be analyzed | `read_list=combined_all_call_hybrids_read_list`<br><br>example list format for paired end data (single end file should contain one line per individual):<br><br>`./reads/CALL1_read1.fq.gz ./reads/CALL1_read2.fq.gz`<br>`./reads/CALL2_read1.fq.gz ./reads/CALL2_read2.fq.gz`<br>`./reads/CALL3_read1.fq.gz ./reads/CALL3_read2.fq.gz` | Always |
| `read_length=` | Provide expected read length | `read_length=150` | Always |
| `prop_genome_genome1_parent=` | Expected proportion of the genome derived from the parent species listed under genome1 | `prop_genome_genome1_parent=0.5` | If not provided, AncestryHMM will attempt to estimate (may increase run time) |
| `number_indiv_per_job=` | Parallelize jobs such that each | `number_indiv_per_job=1` | Always |

| | | | |
|---|---|---|---|
| | job processes this number of individuals.<br><br>Low numbers mean high parallelization and high number mean low parallelization. | | |
| program_path= | Path to the program install folder | program_path=/home/groups/schumer/shared_bin/Ancestry_HMM_pipeline | If not provided the program will assume necessary scripts and programs are in the working directory |
| provide_AIMs= | Coordinates and identities of ancestry informative sites that distinguish the two parent species | provide_AIMs=Xbirchmanni10xgenome_ancestry_informative_sites_filterF1<br><br>Example list format:<br><br>ScyDAA6-2-HRSCAF-26 58345 T C<br>ScyDAA6-2-HRSCAF-26 58976 T A<br>ScyDAA6-2-HRSCAF-26 59896 T C<br>ScyDAA6-2-HRSCAF-26 60164 G A<br>ScyDAA6-2-HRSCAF-26 63105 G A<br>ScyDAA6-2-HRSCAF-26 65532 G A<br>ScyDAA6-2-HRSCAF-26 66290 C A<br>ScyDAA6-2-HRSCAF-26 68233 T C<br>ScyDAA6-2-HRSCAF-26 70398 G A<br>ScyDAA6-2-HRSCAF-26 73869 G A | Required unless provided genomes are on the same coordinate system and can be auto detected |
| provide_counts= | Counts of parental allele frequencies at ancestry informative sites (and recombination rates between adjacent sites if available) | provide_counts=Xbirchmanni10xgenome_Xmalinche_observed_parental_counts_filterF1<br><br>Example format:<br><br>ScyDAA6-2-HRSCAF-26 163722 129 3 0 54 0.00000078<br>ScyDAA6-2-HRSCAF-26 166158 135 5 0 54 0.00001374<br>ScyDAA6-2-HRSCAF-26 166535 6 0 0 6 0.00000754<br><br>Columns are:<br><br>Chromosome site allele1_count_parent1 allele2_count_parent1 allele1_count_parent2 allele2_count_parent2 recombination_rate | If not provided, the program will assume that provided ancestry informative sites are fixed between species |
| per_site_error= | Per-site error parameter for HMM (i.e. due | per_site_error=0.02 | Always |

| | | | |
|---|---|---|---|
| | to sequencing error, contamination, etc) | | |
| gen_initial_admix= | Estimated generation of initial admixture | gen_initial_admix=20 | If not provided, AncestryHMM will attempt to estimate (may increase run time) |
| focal_chrom_list= | Provide a list of chromosomes to run (other chromosomes will not be run) | focal_chrom_list=mychrs.txt<br><br>Example:<br><br>ScyDAA6-2-HRSCAF-26<br>ScyDAA6-7-HRSCAF-50 | Not required |
| rec_M_per_bp= | Estimated recombination rate in Morgans/bp | rec_M_per_bp=0.00000002 | Always; Use an estimate for a related species if not available |
| max_alignments= | Limit analysis to a maximum number of alignments (for computational speed) | max_alignments=2000000 | Optional |
| retain_intermediate_files= | Keep all intermediate files. Warning: setting this to 1 results in a high space footprint for a large run; only recommended for troubleshooting. | retain_intermediate_files=0 | Options are 1 to keep or 0 to delete. |
| posterior_thresh= | Posterior probability threshold to use for identifying | posterior_thresh=0.9 | Recommended 0.8-1 |

| | | | |
|---|---|---|---|
| | ancestry transition intervals | | |
| job_submit_command= | Option to run sequentially if using Docker image for dependencies or from a desktop computer. Set bash to run sequentially and sbatch to run in parallel on a slurm cluter | job_submit_command=bash<br><br>or<br><br>job_submit_command=sbatch | Always required |
| slurm_command_map=<br><br>slurm_command_variant_call=<br><br>slurm_command_hmm= | If running on a slurm cluster, provide cluster specific parameters for queues, time & memory | slurm_command_map=#!/bin/sh #SBATCH --ntasks=1 #SBATCH --cpus-per-task=1 #SBATCH -p schumer --mem=64000 #SBATCH --time=02:30:00<br><br>slurm_command_variant_call=#!/bin/sh #SBATCH --ntasks=1 #SBATCH --cpus-per-task=1 #SBATCH -p schumer --mem=64000 #SBATCH --time=05:00:00<br><br>slurm_command_hmm=#!/bin/sh #SBATCH --ntasks=1 #SBATCH --cpus-per-task=1 #SBATCH -p schumer --mem=64000 #SBATCH --time=03:00:00 | Required if running on a cluster |

**Examples**

Several example files are available with the git repository including example configuration files

**Running the pipeline**

After setting the parameters in the configuration file and loading required dependencies, simply run:

```
perl mixnmatch/ simulate_admixed_genomes_v6.pl
hybrid_simulation_configuration.cfg
```

where path is the path to your simulator install

#### Appendix 3: Protocol used to generate low-coverage sequenced data for $F_1$ and $F_2$ individuals (*X. birchmanni x X. malinche*)

1. **Pre-charge the Tn5 with the adaptors.**

____Combine on ice, ***in the following order***, and mix by pipetting up and down after each addition:

15 µl Tn5 (100 ng / µl)
122 µl reassociation buffer/Glycerol (1:1 mix of reassociation buffer and glycerol, made in advance)
3 µl each adaptor (1 and 2)

____ Incubate in a thermal cycler at 37°C for 30 minutes (with a heated lid).

*Note: this will exceed the allowed volume per well for most thermocyclers, so the mixture should be split between three wells with the appropriate volume per well*

____While the Tn5 is pre-charging, add 3 µl of DNA (3-10 ng /µl) to each well on a 96 well plate (label "tagmentation")

2. **Tagmentation**

____ Make up MasterMix for tagmentation reaction on ice. The volumes per 96 samples (107uL/well mmix in a strip) are:
120 ul Precharged Tn5 from step 1
240 ul 5X TAPS Buffer (contains DMF) – collect waste separately
500 ul H20

____Add 7 uL of the MasterMix to each well of a 96 well plate, already containing your DNA. Mix up and down as you add the MasterMix. Spin briefly.

____Incubate at 55°C for 7 minutes.

3. **Kill the Tn5**

____Add 2.5 µl 0.2% SDS to each reaction, mix up and down, spin briefly.

____Incubate at 55°C for 7 minutes in a thermal cycler.

4. **PCR**

____ Transfer 3 µl of your digested DNA from each well of your old 96 well plate to a new 96 well plate

*Note: Save tagmentation plate in fridge until library concentration and tapestation look good*

____ Add 1 µl of a unique i7 primer from the 96 well i7 plate (10 µM i7) to each well of this plate.

____ Make up master mix for PCR for 96 samples (171uL/well mmix in a strip)
120 µl 10 µM i5 primer
900 µl OneTaq HS Quick-Load 2X
350 µl H20

____ Add 11 µl of the MasterMix to each well of the new 96 well plate. Mix by pipetting up and down as you add the MasterMix. Spin plate briefly.

Run on thermal cycler with following protocol:

| Step | Temp | Time |
|---|---|---|
| 1 | 68°C | 3 min |
| 2 | 95°C | 30 sec |
| 3 | 95°C | 10 sec |
| 4 | 55°C | 30 sec |
| 5 | 68°C | 30 sec |
| 6 | Cycle to step 3 less than 12 times **Note: the fewer cycles, the better the sequencing results** | |
| 7 | 68°C | 5 minutes (final extension) |
| 8 | 4°C | Hold |

#### 5. Clean up library

___ Pipette 5-10 µl from each well of the plate into a reservoir or set of strip tubes, and collect this volume into a 2 mL Eppendorf tube. For a full plate with 10 µl collected from each well, make two 400 µl aliquots

___ Remove bottle of 18% SPRI beads from fridge, and allow beads to come to room temperature.

___ Gently shake bottle to resuspend beads (make sure no beads remain settled at the bottom of the bottle).

___ Add 1 volume of beads, mix by pipetting up and down 10 times.

___ Incubate at room temperature for 5 minutes.

___ Place on magnet for 5 minutes.

___ Aspirate and discard supernatant. Collect in DMF waste container.

___ Add 500 µl of ETOH 70%, wash the sides of the Eppendorf tube while adding

___ Incubate at room temperature for 1 minute.

___ Aspirate and discard supernatant.

___ Add 500 µl of ETOH 70% again, wash the sides of the Eppendorf tube while adding

___ Incubate at RT 1 min.

___ Aspirate and discard supernatant. Remove as much ethanol as possible without disturbing the magnetic ball/ring.

___ Air dry 5 minutes.

___ Off the magnet, resuspend in 17 µl of Qiagen Buffer EB.

___ Incubate at room temperature for 3 minutes.

___ Place back on magnet for 5 minutes.

___ Transfer the supernatant (15 µl) to a new tube.

6. **Qubit Library**

7. **Tape station or bioanalyze your library**

**Instructions on how to make buffers and adapter stocks for Tn5 libraries:**

**Adapter 1 & 2 complex:**

Adapter 1 (10 uM)
10uL Tn5ME-A (100 uM)
10uL Tn5MErev (100 uM)
80uL Reassociation Buffer

Adapter 2 (10 uM)
10uL Tn5ME-B (100 uM)
10uL Tn5MErev (100 uM)
80uL Reassociation Buffer

____ Mix well.

____ Anneal the oligonucleotides that will form the free-end adaptors in a thermal cycler with the following Anneal Program:

| Step | Temp | Time |
|---|---|---|
| 1 | 95°C | 10 min |

| | | |
|---|---|---|
| 2 | 90°C | 1 min |
| 3 | Reduce temp by 1°C/cycle 60 times in thermocycler | |
| 4 | 4°C | Hold |

**Reassociation Buffer:**
10 mM Tris pH 8.0
50 mM NaCl
1 mM EDTA

**TAPS buffer:**
*– store at -20 for long time storage and at 4C for a few weeks, avoid freeze thaw cycles.*
5x TAPS-DMF buffer from Picelli paper:
50 mM TAPS-NaOH,
25 mM MgCl2,
50% v/v DMF (pH 8.5) at 25°C

**Adapter sequences:**

Tn5MErev: 5’-[phos]CTG TCTCTTATACACATCT-3’;
Tn5ME-A: 5’- TCGTCGGCAGCGTCAGATGTGTATAAGAGACAG-3’
Tn5ME-B: 5’-GTCTCGTGGGCTCGGAGATGTGTA TAAGAGACAG-3’.

**i5:** AAT GAT ACG GCG ACC ACC GAG ATC TAC AC NNNNNNNN[+] TC GTC GGC AGC GTC

[+]plate barcode

**i7:** CAA GCA GAA GAC GGC ATA CGA GAT NNNNNNNN[+] G TCT CGT GGG CTC GG

[+]96 different sequences for sample barcode

#### Appendix 4: Example configuration files for specific simulations

*Base simulation:*

```
genome1=ScyDAA6-1508-HRSCAF-1794_section.fa
genome2=
use_ancestral=1
mixture_prop_par1=0.5
rec_rate_Morgans_kb=0.00002
num_indivs=50
gens_since_admixture=200
chr_to_simulate=group1
poly_perbp_par1=0.001
poly_perbp_par2=0.001
rate_shared_poly_at_aims=
read_type=PE
read_length=100
per_bp_indels=0.006
sequencing_error=0.005
number_reads=25000
parental_drift=0
macs_par1_aims_pop=
macs_par2_aims_pop=
aim_freq_cutoff=0.95
cross_contam=
job_submit_cmd=sbatch
use_macs=1
use_map=1
SELAM_param_file=
SELAM_selection_file=
macs_params=200 10000000 -I 2 100 100 0 -t 0.001 -h 1e2 -r 0.001 -ej 2 2 1 -R
recombination_map_for_macs_section.txt
program_path=/scratch/groups/schumer/molly/Simulate_Ancestry_HMM/Simulate_hybrid_genomes
par1_for_aims=20
par2_for_aims=20
seq_params=
job_header=#!/bin/sh #SBATCH --ntasks=1 #SBATCH --cpus-per-task=1 --mem=64000 #SBATCH -p
schumer #SBATCH --time=02:00:00
num_indiv_per_job=2
```

*Source population drift:*

```
genome1=ScyDAA6-1508-HRSCAF-1794_section.fa
genome2=
use_ancestral=1
mixture_prop_par1=0.5
rec_rate_Morgans_kb=0.00002
num_indivs=50
gens_since_admixture=200
chr_to_simulate=group1
poly_perbp_par1=0.001
poly_perbp_par2=0.001
rate_shared_poly_at_aims=0.01
read_type=PE
read_length=100
per_bp_indels=0.006
sequencing_error=0.005
number_reads=100000
parental_drift=1
macs_par1_aims_pop=3
macs_par2_aims_pop=4
aim_freq_cutoff=0.9
cross_contam=
job_submit_cmd=sbatch
use_macs=1
use_map=1
SELAM_param_file=
SELAM_selection_file=
macs_params=220 10000000 -I 4 100 100 20 20 0 -t 0.001 -h 1e2 -r 0.001  -ej 0.5 3 1 -ej 0.51 4
2 -ej 2 2 1 -R recombination_map_for_macs_section.txt
program_path=/scratch/groups/schumer/molly/Simulate_Ancestry_HMM/Simulate_hybrid_genomes
par1_for_aims=20
par2_for_aims=20
seq_params=
job_header=#!/bin/sh #SBATCH --ntasks=1 #SBATCH --cpus-per-task=1 --mem=64000 #SBATCH -p
schumer #SBATCH --time=02:00:00
num_indiv_per_job=1
```

*Historical migration between parental populations:*

```
genome1=ScyDAA6-1508-HRSCAF-1794_section.fa
genome2=
use_ancestral=1
mixture_prop_par1=0.5
rec_rate_Morgans_kb=0.00002
num_indivs=50
gens_since_admixture=200
chr_to_simulate=group1
poly_perbp_par1=0.001
poly_perbp_par2=0.001
rate_shared_poly_at_aims=
read_type=PE
read_length=100
per_bp_indels=0.006
sequencing_error=0.005
number_reads=25000
parental_drift=0
macs_par1_aims_pop=
macs_par2_aims_pop=
aim_freq_cutoff=0.95
cross_contam=
job_submit_cmd=sbatch
use_macs=1
use_map=1
SELAM_param_file=
SELAM_selection_file=
macs_params=200 10000000 -I 2 100 100 0.025 -t 0.001 -h 1e2 -r 0.001 -ej 2 2 1 -R
recombination_map_for_macs_section.txt
program_path=/scratch/groups/schumer/molly/Simulate_Ancestry_HMM/Simulate_hybrid_genomes
par1_for_aims=20
par2_for_aims=20
seq_params=
job_header=#!/bin/sh #SBATCH --ntasks=1 #SBATCH --cpus-per-task=1 --mem=64000 #SBATCH -p
schumer #SBATCH --time=02:00:00
num_indiv_per_job=2
```

*Hybridization with selection:*

```
genome1=ScyDAA6-1508-HRSCAF-1794_section.fa
genome2=
use_ancestral=1
mixture_prop_par1=0.5
rec_rate_Morgans_kb=0.00002
num_indivs=50
gens_since_admixture=200
chr_to_simulate=group1
poly_perbp_par1=0.001
poly_perbp_par2=0.001
rate_shared_poly_at_aims=
read_type=PE
read_length=100
per_bp_indels=0.006
sequencing_error=0.005
number_reads=25000
parental_drift=0
macs_par1_aims_pop=
macs_par2_aims_pop=
aim_freq_cutoff=0.95
cross_contam=
job_submit_cmd=sbatch
use_macs=1
use_map=1
SELAM_param_file=
SELAM_selection_file=SELAM_selection_file
macs_params=200 10000000 -I 2 100 100 0 -t 0.001 -h 1e2 -r 0.001 -ej 2 2 1 -R
recombination_map_for_macs_section.txt
program_path=/scratch/groups/schumer/molly/Simulate_Ancestry_HMM/Simulate_hybrid_genomes
par1_for_aims=20
par2_for_aims=20
seq_params=
job_header=#!/bin/sh #SBATCH --ntasks=1 #SBATCH --cpus-per-task=1 --mem=64000 #SBATCH -p
schumer #SBATCH --time=02:00:00
num_indiv_per_job=2
```

*associated SELAM selection file:*

```
# cat selection_file

S	A	0	0.1	1	1	0.75
```

*Simulating multiple hybridization pulses:*

```
genome1=ScyDAA6-1508-HRSCAF-1794_section.fa
genome2=
use_ancestral=1
mixture_prop_par1=0.5
rec_rate_Morgans_kb=0.00002
num_indivs=50
gens_since_admixture=2000
chr_to_simulate=group1
poly_perbp_par1=0.001
poly_perbp_par2=0.001
rate_shared_poly_at_aims=
read_type=PE
read_length=100
per_bp_indels=0.006
sequencing_error=0.005
number_reads=25000
parental_drift=0
macs_par1_aims_pop=
macs_par2_aims_pop=
aim_freq_cutoff=0.95
cross_contam=
job_submit_cmd=sbatch
use_macs=1
use_map=1
SELAM_param_file=selam_demography.txt
SELAM_selection_file=
macs_params=200 10000000 -I 2 100 100 0 -t 0.001 -h 1e2 -r 0.001 -ej 2 2 1 -R
recombination_map_for_macs_section.txt
program_path=/scratch/groups/schumer/molly/Simulate_Ancestry_HMM/Simulate_hybrid_genomes
par1_for_aims=20
par2_for_aims=20
seq_params=
job_header=#!/bin/sh #SBATCH --ntasks=1 #SBATCH --cpus-per-task=1 --mem=64000 #SBATCH -p
schumer #SBATCH --time=02:00:00
num_indiv_per_job=2
```

*associated SELAM demography file:*

```
# cat selam_demography.txt

Pop1   Pop2   Sex    0      1      1800   1801
0      0      A      5000   5000   5000   5000
0      a0     A      0.5    0      0.25   0
0      a1     A      0.5    0      0.25   0
```
